# The mitochondrial disulphide relay substrate FAM136A safeguards IMS proteostasis and cellular fitness

**DOI:** 10.1101/2025.09.22.677734

**Authors:** Christine Zarges, Hanna Fieler, Robin Alexander Rothemann, Simon Poepsel, Lucas T Jae, Jan Riemer

## Abstract

The mitochondrial disulphide relay is the key machinery for import and oxidative protein folding in the mitochondrial intermembrane space. Among IMS proteins with unknown function, we identified FAM136A as a new substrate of the mitochondrial disulphide relay. We demonstrate a transient interaction between FAM136A and MIA40, and that MIA40 introduces four disulphide bonds in two twin-CX_3_C motifs of FAM136A. Consequently, IMS import of FAM136A requires these cysteines and its steady state levels in intact cells are strongly dependent on MIA40 and AIFM1 levels. Furthermore, we show that FAM136A forms non-covalent homodimers as a mature protein. Acute deletion of FAM136A curtails cellular proliferation capacity and elicits a robust induction of the integrated stress response, coincident with the aggregation and/or depletion of selected IMS proteins including HAX1 and CLPB. Together, this establishes FAM136A as a pivotal component of the IMS proteostasis network, with implications for overall cellular function and health.

## INTRODUCTION

The mitochondrial intermembrane space (IMS) houses a comparably small, specialized subset of less than 10% of the mitochondrial proteome [1–3]. However, IMS proteins are involved in critical functions such as regulation of respiration, metabolism, and coordination of mitochondrial activities with other cellular compartments [4, 5]. Many IMS proteins participate in biogenesis and maintenance of functional respiratory chain complexes. Others are involved in protein import, folding and degradation. Some IMS proteins also play important roles in apoptosis initiation and regulation. Further, the unique location of the IMS, sandwiched between the outer (OMM) and inner (IMM) mitochondrial membranes, allows its proteins to act as intermediaries in various signalling and metabolic pathways, making the IMS essential for overall mitochondrial function and cellular homeostasis. Disruptions in IMS proteostasis, such as the accumulation of misfolded or damaged proteins, can activate stress signalling pathways, including the integrated stress response (ISR) [6, 7]. Among other things, the ISR leads to a reduction of global protein synthesis while selectively enhancing the translation of stress-adaptive factors, such as ATF4.

Interestingly, this pathway can be detrimental [8, 9], but has also proven cytoprotective in certain genetic contexts [10–12].

Notably, the majority of IMS proteins contain conserved cysteines, which play crucial roles in protein structure and function [1, 13–17]. These cysteines contribute to protein stability through disulphide bond formation, and provide (reactive) thiol groups for catalysis, metal binding, or redox sensing. Mutations in specific cysteines of IMS proteins have been associated with different pathologies. For example, mutations in TIMM8A (C66W) and NDUFB10 (C107S) have been reported to cause infantile mitochondrial disorders [18, 19].

In the IMS, the mitochondrial disulphide relay constitutes the key machinery for oxidative protein folding [1, 13–17]. In human cells, it consists of three core components, the oxidoreductase MIA40 (also CHCHD4), the sulfhydryl oxidase ALR, and AIFM1, which is important for MIA40 import and appears to have specialized MIA40-activating functions within the machinery [20–23]. MIA40 contains two key features, a redox-active cysteine motif formed by cysteines 53 and 55 (“CPC”) and a hydrophobic groove that enables MIA40 to function as a chaperone holdase [24–28]. The redox-active cysteines of MIA40 are usually present in their oxidized state which allows them to form mixed disulphide bonds with reduced substrate proteins once they traverse the OMM via the translocase of the outer membrane (TOM) [29]. A prerequisite for this redox reaction is the correct orientation of the substrate which is facilitated through binding to the hydrophobic groove of MIA40 [24]. As a result of the interaction with MIA40, substrates acquire intramolecular disulphide bonds which fold and stabilize them. Notably, as most classical MIA40 substrates do not contain mitochondrial targeting signals, disulphide formation is also coupled to their import and thereby traps folded substrates in the IMS [27, 30–33]. After substrate oxidation, MIA40 is reduced and subsequently re-oxidized by ALR. ALR transfers the electrons to cytochrome *c* and ultimately to complex IV of the electron transport chain, completing the relay [26, 34–37].

When screening the orphan IMS proteome [3] for proteins with conserved cysteines, we identified FAM136A. FAM136A has been genetically associated with Menier’s disease [38, 39], a chronic inner ear disorder characterized by recurring episodes of vertigo, fluctuating hearing loss, tinnitus, and aural fullness. A nonsense mutation in FAM136A introducing a premature stop codon disrupted the FAM136A protein product and led to significantly decreased expression of FAM136A transcripts in carriers’ lymphoblastoid cell lines [39]. Notably, this nonsense variant has only been reported in one family, and GnomAD population data shows that in principle loss of function variants in FAM136A are tolerated [40]. *In silico* analysis of existing genomic data using the DepMap and Fireworks databases [41] revealed close functional relationships between FAM136A and genes encoding for HTRA2, CLPB and MTCH2. CLPB and HTRA2 have been described as part of the IMS proteostasis network [42, 43] and MTCH2 takes a crucial role in OMM protein biogenesis [44]. Thus, FAM136A might play a role in IMS proteostasis and biogenesis of certain IMS and OMM proteins. Still, the pathological mechanisms of FAM136A mutations and the physiological functions of FAM136A remain poorly understood. FAM136A has recently been found localized to the IMS in a crosslinking proteome study and subsequent biochemical experiments confirmed this localization [45]. Moreover, FAM136A also emerged as hit in a genetic screen for growth in galactose-based media, hinting at a potential role in supporting OXPHOS [46]. Intriguingly, loss of FAM136A led to destabilization of selected IMS proteins including HAX1 and ENDOG [46] and the induction of a mitochondrial unfolded protein response [47] indicating a role of the protein in IMS proteostasis. Notably, FAM136A also appeared repeatedly in MIA40 interactome studies indicating its potential to form disulphide bonds [48]. We hypothesized that FAM136A could be a new disulphide relay substrate and investigated FAM136A biogenesis. This revealed that it indeed constitutes a novel client of this axis, thereby expanding the substrate spectrum of this critical import pathway. Additionally, we find that defects in FAM136A reduce cellular growth and trigger the ISR, highlighting an important role of FAM136A in maintaining IMS proteostasis and cellular fitness.

## RESULTS

### FAM136A – a soluble orphan IMS protein with conserved cysteines

The IMS contains about 96 proteins according to MitoCarta 3.0 and different proteomics data sets [3, 5, 13, 49], including 21 proteins without any known function and unclear biogenesis. Among these, we identified conserved cysteines across the model organisms, *M. musculus*, *D. rerio*, *X. laevis*, *D. melanogaster*, and *C. elegans* in 15 proteins (**Figure 1A**). The protein FAM136A was of particular interest as it contains nine cysteine residues of which eight are present in conserved CX_3_C motifs – hallmarks of the family of small TIM chaperones (**Figure 1B**, *always 4 cysteines forming one twin-CX_3_C motif*, [50]). The structural prediction for FAM136A shows a bundle of three alpha-helical segments, the interaction of which is stabilized by these CX_3_C mediated disulphide bridges. Indeed, the C-terminal part of FAM136A (aa60-138) shows strong sequence homology to the TIM chaperone TIM10 for which the arrangement and stabilization of α-helices via CX_3_C motifs has been experimentally shown [51]. The IMS localization of FAM136A has been demonstrated in an APEX-based IMS proteome [3] and in a crosslinking proteome that also demonstrated its localization biochemically [45]. We confirmed these findings by performing cellular fractionation, immunofluorescence microscopy, mitochondrial subfractionation and a carbonate extraction assay in HEK293 cells (**Figure 1C-F**). Thus, FAM136A is an uncharacterized soluble IMS protein with eight conserved cysteines and no classical mitochondrial targeting signal.

**Figure 1.**
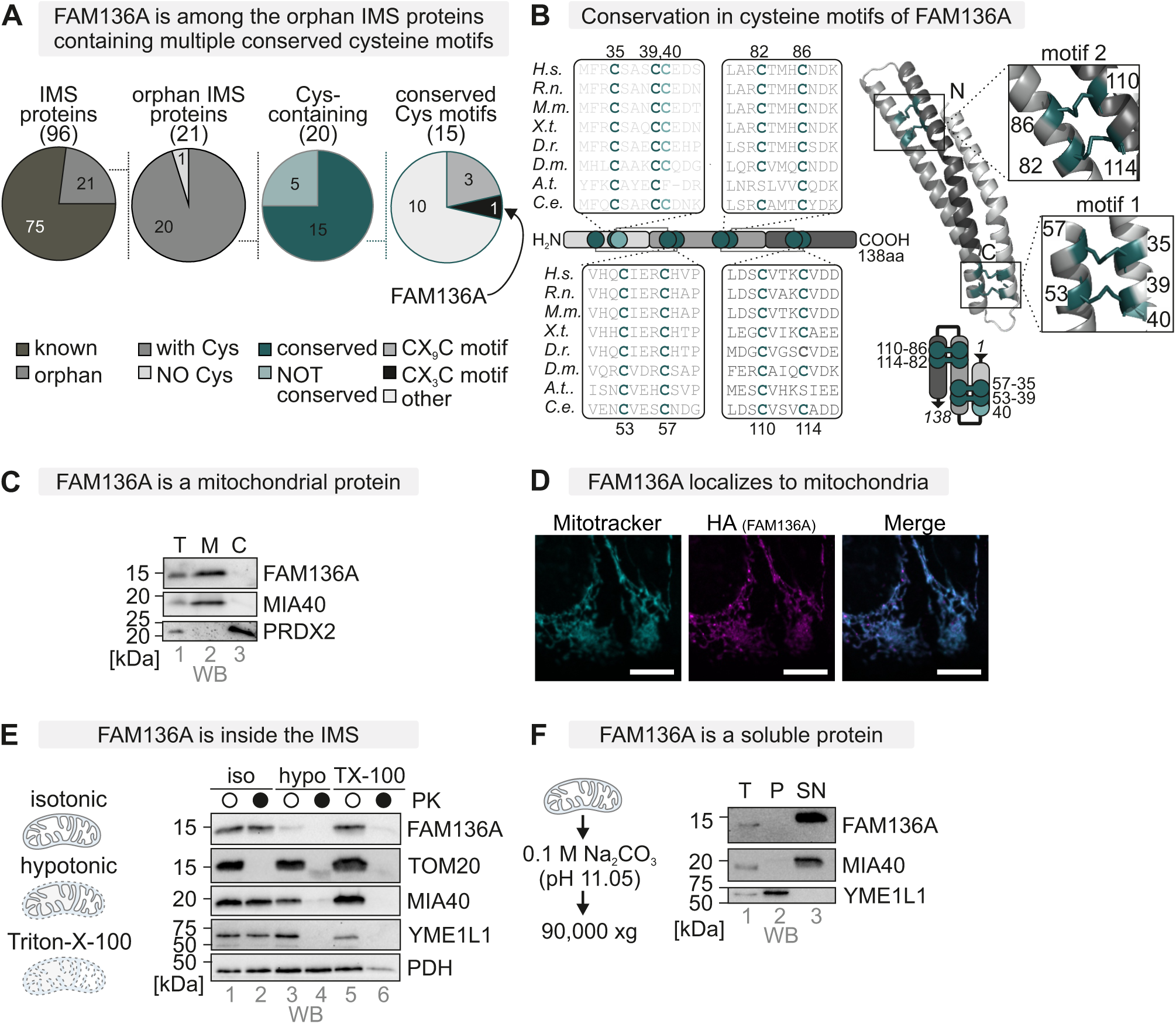
FAM136A is a soluble IMS protein with conserved cysteines **A.** *In silico* analysis of proximity-labeling data and IMS proteomics data [3, 5, 13, 49]. Of the 96 IMS proteins (soluble or with domains in the IMS, 21 are not fully characterized. Of those, 20 contain cysteines. In 15 of the protein’s cysteines are conserved. In those 15 proteins, the cysteines are positioned in three in twin-CX_9_C and in one in a twin-CX_3_C motif. **B.** Conserved cysteines across species and Alpha-fold3 prediction. H.s., Homo sapiens, R.n., Rattus norvegicus, M.m., Mus musculus, X.t., Xenopus tropicalis, D.r., Dario rerio, D.m., Drosophila melanogaster, A.t., Arabidopsis thaliana, C.e., Caenorhabditis elegans. Cysteines are highlighted and the boxes with varying grey colours indicate the helices predicted for FAM136A. **C.** Localization of FAM136A by cellular fractionation. Cells were lysed by potter homogenization followed by centrifugation to separate a mitochondrial (“M”) and a post-mitochondrial (“C”) fraction. The total lysate (“T”) served as loading control, and MIA40 (arrowhead) and PRDX2 as controls for mitochondria and cytosol, respectively. **D.** Localization of FAM136A by immunofluorescence microscopy. Cells were fixated, permeabilized, and stained using a primary antibody against the HA-tag or mitotracker. Cells were analysed by fluorescence microscopy. Mitotracker signal served as positive control for mitochondria. FAM136A-HA localizes to mitochondria in HEK293 cells. Magenta, HA signal; Cyan, mitotracker signal. A lilac/whitish colour indicates signal overlap. Bar corresponds to 10 μm. **E.** Localization of FAM136A by mitochondrial fractionation. Mitochondria were enriched as described in (C.) and then either incubated in an isotonic buffer (“iso”), in a hypotonic buffer (“hypo”) or a buffer containing triton X-100 (TX-100”). The resulting fraction were either treated with proteinase K (“PK”) to test for protection of proteins or left untreated. FAM136A is protected in isolated mitochondria (*lane 2*) but is lost already after hypotonic treatment of mitochondria indicating it to be a soluble IMS protein. TOM20, MIA40, YME1L and PDH served as OMM, IMS, IMM and matrix controls, respectively. **F.** Solubility of FAM136A by carbonate extraction assay. Isolated mitochondria were treated with Na_2_CO_3_ and then supernatant and pellet were separated by ultracentrifugation. FAM136A behaves like the soluble control MIA40 and not like the membrane control YME1L.

### FAM136A undergoes oxidation by MIA40 during its maturation

We next tested whether the cysteines of endogenous FAM136A were indeed present in their oxidized state in intact cells. To this end, we employed a direct redox shift assay (**Figure 2A**). This assay relies on the accessibility of reduced, but not oxidized, thiol residues to the alkylating reagent methyl-polyethylenglycol-maleimide (mmPEG_12_) that causes a mass increase of the modified protein (**Figure 2B**, *lane 4 [reduced] and 2 [oxidized]*). Using this assay, we found that at steady state virtually all FAM136A was present in the oxidized state (**Figure 2B**, *lane 3*).

**Figure 2.**
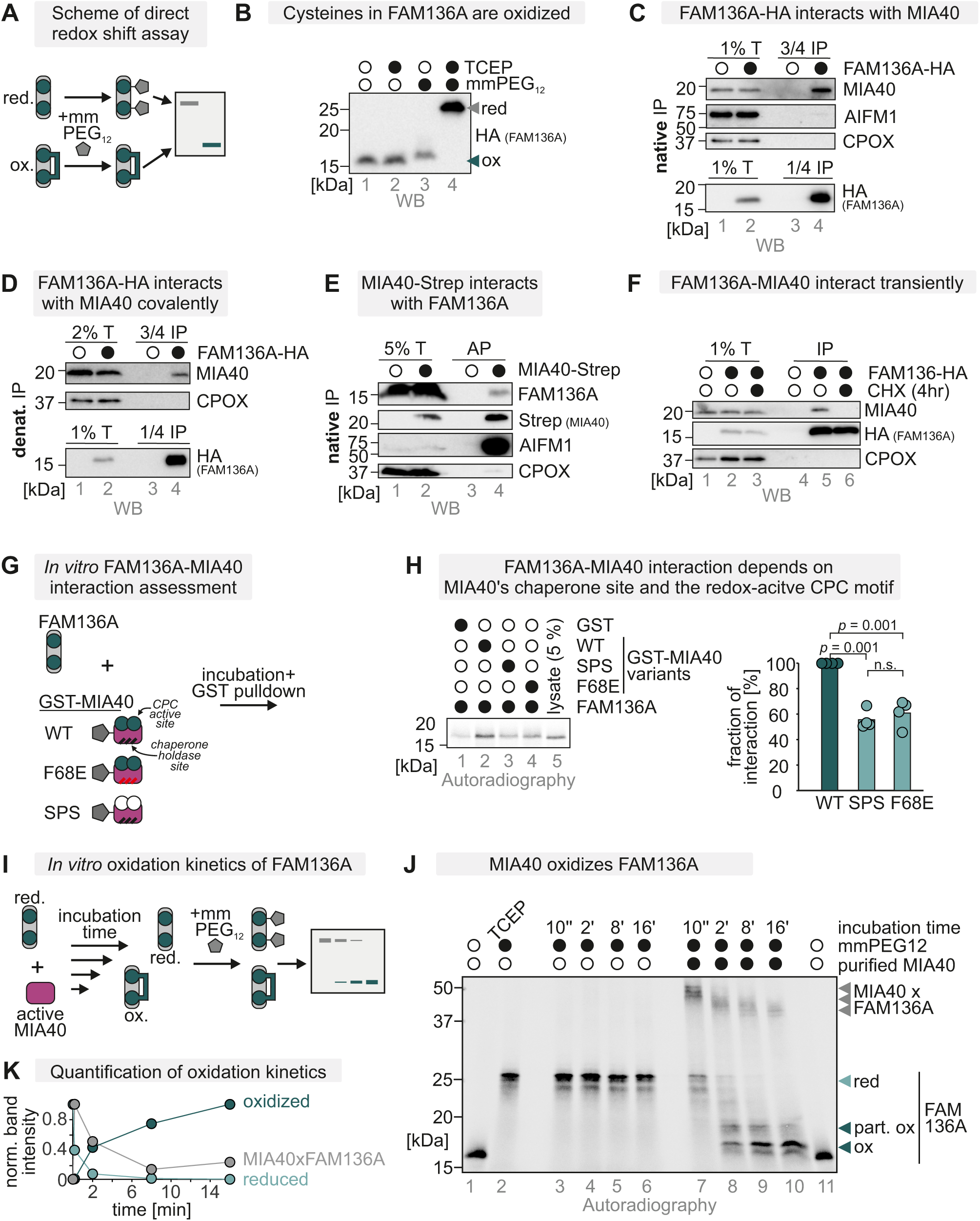
FAM136A becomes oxidized by MIA40 during its maturation **A.** Direct redox shift assay to assess redox state of FAM136A. Cells were lysed and treated with the maleimide mmPEG_12_ that modifies free thiols (reduced, red.) but not thiols in disulfide bonds (oxidized, ox.). Modification of proteins with mmPEG_12_ results in a slower migration of the protein on SDS-PAGE. **B.** Redox state analysis of FAM136A as described in (A.). To test for the redox state of FAM136A, cells were lysed and either treated with the strong reductant TCEP (lanes 2 and 4) or left untreated (lane 1 and 3). Then lysates were left untreated (lanes 1 and 2) or incubated with mmPEG_12_ (lanes 3 and 4). Lysates were analyzed by SDS-PAGE and immunoblotting. Signals were quantified using Image Lab, and the amount of reduced and oxidized protein was plotted. FAM136A-HA is mainly present in the oxidized state. Virtually all of the protein is oxidized at steady state. Black and white circles represent the indicated treatment and not-treated samples. N = 3 replicates. **C.-E.** Assessment of the MIA40-FAM136A interaction. FAM136A-HA (C.,D.) or MIA40-Step (E) were precipitated (immunoprecipitation with HA-beads [IP] or affinity precipitation using streptactin beads [AP]) after stopping thiol-disulfide exchange reactions by NEM incubation from HEK293 cells stably and inducibly expressing these proteins. Precipitates were tested for MIA40 (C.,D.) or FAM136A (E.). Indicated amounts of the total lysate were loaded as input control. FAM136A coprecipitates MIA40 and *vice versa*. For the FAM136A-HA IP, this coprecipitation was also present when cells were lysed under denaturing conditions (D.) indicating a covalent interaction between both proteins. In (C.-E.), CPOX served as negative control, while in (E.), AIFM1 served as positive control. Black and white circles represent the presence of the indicated protein. **F.** Duration of the FAM136A–MIA40 interaction. HEK293 cells stably expressing FAM136A-HA were treated for 4 hours with the translation inhibitor cycloheximide (CHX). Then, cells were lysed under native conditions, and FAM136A-HA was precipitated. Precipitates were analysed by reducing SDS– PAGE and immunoblotting. MIA40 and FAM136A interact transiently with each other comparable to other MIA40 substrates. CPOX served as negative control. **G.** Scheme of assay to assess MIA40-FAM136A interaction *in vitro*. GST-tagged variants of MIA40 (wild-type, WT, redox-inactive C4, 53, 55S (SPS) and chaperone-inactive F68E (FE) and GST alone were purified and bound to beads. The beads were incubated with *in vitro* translated radioactive FAM136A. Subsequently, beads were washed and analysed for bound FAM136A by SDS-PAGE and autoradiography. Incubation with GST-only bound beads served as control. **H.** *In vitro* assessment of MIA40-FAM136A interaction. WT MIA40 (lane 2) interacts with FAM136A. By comparison, the interaction between FAM136A and the MIA40 C4, 53, 53S (lane 3) or the MIA40 F68E variants (lane 4) were significantly decreased (one-way ANOVA with post hoc Tukey HSD test). The control (GST-only bound beads) revealed only minor background binding (lane 1). 5% of the total lysate was loaded as input control (lane 5). TCE staining served as loading control. For quantification, the co-precipitated FAM136A levels were normalized to the levels precipitated by WT MIA40 and in addition normalized to the according MIA40 signal in the TCE staining. Black and white circles represent the indicated treatment and not-treated samples. N = 3 **I.** *In vitro* oxidation kinetics assay of FAM136A. Recombinant expressed and purified human MIA40 WT was added to ^35^S-labeled FAM136A to allow disulphide exchange reactions. The reaction was stopped by rapid acidification via addition of trichloroacetic acid (TCA). Lysates were treated with mmPEG_12_ to determine protein redox states, followed by non-reducing SDS-PAGE and autoradiography. **J.** Assessment of FAM136A oxidation by purified MIA40 as described in (I). Reduced FAM136A is modified with nine mmPEG_12_, whereas oxidized FAM136A was modified by only one mmPEG_24_. In the control experiment, FAM136A did not become oxidized in the absence of MIA40 during the indicated times. Addition of MIA40 to FAM136A resulted in an oxidation of the protein over time. Reduced and oxidized forms as well as semi-oxidized intermediates of FAM136A, and the FAM136A-MIA40 mixed disulphide complex are indicated. N = 3 replicates.

The presence of disulphide bonds indicates that FAM136A might interact with the mitochondrial disulphide relay, specifically its oxidoreductase MIA40. Indeed, in previous high throughput experiments, FAM136A was identified as a possible MIA40 interaction partner [48]. To verify this, we performed targeted co-immunoprecipitation experiments and found that FAM136A co-precipitated with MIA40 and *vice versa* under native (**Figures 2C,E S1A,B**) and denaturing (**Figures 2D)** conditions, implying a covalent interaction between both proteins. For other disulphide relay substrates, it has been demonstrated that they only interact with MIA40 during import and maturation. The interaction between substrates and MIA40 was lost in experiments in which cells were treated with translation inhibitors because no new mitochondrial precursors entered the IMS [22, 52, 53]. In line with this, we found a similar transient nature of the MIA40–FAM136A interaction in a cycloheximide chase treatment-coupled immunoprecipitation experiment (**Figure 2F**).

To further investigate the nature of this interaction, we turned to an *in vitro* reconstituted system. We purified MIA40 wild-type (MIA40 WT), a MIA40 variant that lacks the redox-active cysteine residues C53 and C55 (MIA40 SPS), and a variant impaired in the chaperone function (MIA40 F68E). We coupled the GST-tagged versions of these MIA40 variants to beads and incubated them with [^35^S]-labelled *in vitro* translated FAM136A-HA (**Figure 2G**). Subsequent pelleting of the beads revealed that FAM136A interacted with the WT and to a significantly lesser extent with the redox-inactive and the chaperone-impaired MIA40 variants (**Figure 2H**, *lanes 2-4*). All variants interacted better with FAM136A than the GST negative control. This indicates that both the oxidoreductase and the chaperone function of MIA40 are involved in the interaction with FAM136A.

Lastly, we explored whether FAM136A could be oxidized by MIA40 *in vitro* (**Figures 2I-K**). We incubated an excess of purified oxidized MIA40 WT with [^35^S]-labelled, *in vitro* translated reduced FAM136A-HA. The use of excess MIA40 excluded the necessity of reoxidation of MIA40 during the assay. Upon mixing FAM136A and MIA40, a disulphide-linked complex between MIA40 and FAM136A was rapidly generated (**Figure 2J**, *lane 7*). This FAM136A-MIA40 complex represented an intermediate, which disappeared concomitantly with the appearance of oxidized forms of monomeric FAM136A (**Figure 2J**, *lanes 8-11, 8 out of the 9 cysteines of FAM136A seem to be oxidized*). The generation of a fully oxidized form demonstrated that MIA40 alone was sufficient to generate all disulphide bonds in FAM136A (**Figure 2J**, *arrowhead*). Moreover, various semi-oxidized FAM136A species and FAM136A-MIA40 complexes in which FAM136A already contained internal disulphide bonds appeared and persisted throughout the incubation time (**Figure 2J**, *lanes 7-11*). Without the addition of MIA40, FAM136A remained reduced throughout (**Figure 2J**, *lanes 3-6*).

In sum, FAM136A transiently interacts with MIA40 under formation of transient mixed MIA40-FAM136A disulphide bonds which become dissolved by introducing four disulphide bonds into FAM136A.

### FAM136A IMS import and stability depend on the mitochondrial disulphide relay

To test whether FAM136A relies on MIA40 for its import, we performed *in organello* import experiments in mitochondria isolated from WT, AIFM1 knockout (KO) and MIA40 knockdown (MIA40 KD) HEK293 cells (**Figures 3A-C**) [54]. Isolated mitochondria were incubated with [^35^S]-labelled, *in vitro* translated FAM136A-HA protein lysate and after specific times, all non-imported proteins were degraded by proteinase K treatment (**Figure 3A**). FAM136A could be imported well into WT mitochondria, but its import was strongly and significantly affected by loss of MIA40 or AIFM1 (**Figures 3B,C**). In addition, its import was also depended on the mitochondrial membrane potential, as treatment with the IMM depolarizing agent carbonyl cyanide 3-chlorophenylhydrazone (CCCP), ablated import (**Figures 3B,C**, *lanes 5,9*). The import of the established MIA40 substrate COX19 [32] into these mitochondria was hampered to a similar extent.

**Figure 3.**
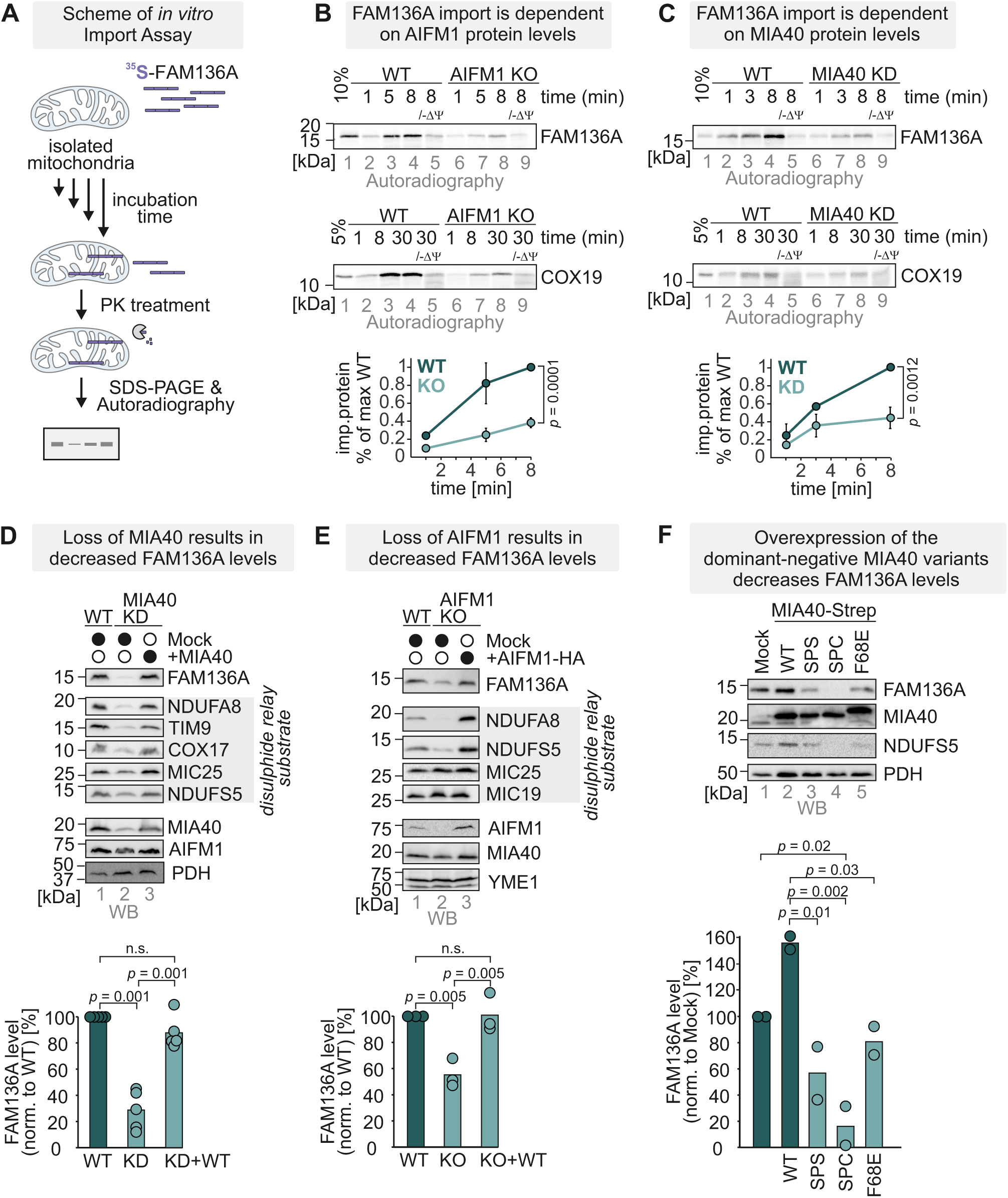
IMS import of FAM136A depends on the mitochondrial disulphide relay **A-C.** *In organello* import assays with FAM136A. *In vitro* translated radioactive FAM136A was incubated with mitochondria isolated either from MIA40 knockdown (KD, B) or AIFM1 knockout (KO, C) HEK293 cells. Non-imported proteins were removed by treatment with Proteinase K. An import reaction was performed on mitochondria treated with CCCP to dissipate the mitochondrial membrane potential (–ΔΨ). Imported proteins were analyzed by reducing SDS-PAGE and autoradiography. Signals were quantified using ImageQuantTL, and the amount of imported protein was plotted. FAM136A import is strongly dependent on both MIA40 and AIFM1 (one-sample t-test). In addition, it relies on the mitochondrial membrane potential for import. Import of COX19 served as positive control. N = 3 replicates. **D,E.** Levels of endogenous FAM136A in cells lacking MIA40 (MIA40 KD) (D) or AIFM1 (AIFM1 KO) (E). Protein levels in HEK293 cell lines depleted of MIA40 (D) or AIFM1 (E) or overexpressing the respective protein were compared to protein levels in wild-type HEK293 cells. Lysates were analysed by reducing SDS-PAGE and immunoblotting. Signals were quantified using ImageLab, and the amount of protein was plotted. TCE staining served as loading control. FAM136A levels were decreased in cells lacking MIA40 or AIFM1 significantly (one-way ANOVA with post hoc Tukey HSD test). Black and white circles represent the indicated treatment and not-treated samples. N = 5 (D) or 3 (E) replicates **F.** Steady-state levels of FAM136A in HEK293 cells stably expressing different variants of MIA40. Expression of MIA40 variants lacking either both cysteines of the redox active CPC motif (SPS) or only C53 (SPC) or with a mutation in the chaperone site of MIA40 (F68E) was induced for 5 days in glucose-containing medium. Cells were lysed, and protein levels were analyzed by SDS-PAGE and immunoblot against the indicated proteins. Overexpression of MIA40 variants decreases FAM136A steady state levels. (one-way ANOVA with post hoc Tukey HSD test). N = 2

We next investigated the levels of FAM136A in MIA40 KD and AIFM1 KO cells and found that FAM136A levels were reduced to about 30-50% in both cell lines, well in line with other substrates of MIA40 (**Figures 3D,E**). These values are also in accord with high-throughput proteomics studies that indicated a reduction of FAM136A levels to 17% [48] and 27% [22] in MIA40 KD and AIFM1 KO cells, respectively.

Interestingly, overexpression of MIA40 in the MIA40 KD background restored FAM136A back to WT levels but did not lead to even higher protein levels of FAM136A (**Figure 3D**, *lane 3*). For other MIA40 substrates, including COX17 and NDUFA8, such a correlation between MIA40 and substrate levels was observed, indicating that MIA40 levels are limiting for the handling of those proteins but not for FAM136A [28, 32, 48]. We also tested FAM136A levels in cells overexpressing different MIA40 variants: the MIA40 SPS variant lacking the redox active CPC motif, the SPC variant lacking cysteine C53 of this motif, and the F68E variant bearing a mutation in the chaperone site of MIA40. These variants, in particular the SPC variant, have previously been demonstrated to impact MIA40 substrate import and levels negatively [32, 48]. In line with this, we also found strongly reduced levels of FAM136A upon overexpression of any of these variants for extended times (**Figure 3F**). Thus, the disulphide relay is critical for mitochondrial accumulation and stability of FAM136A.

### FAM136A cysteines determine its stability

FAM136A cysteines become oxidized by the disulphide relay during import. We thus tested their importance for cellular localization and *in organello* protein import and stability. To this end, we constructed a series of cell lines that express different cysteine variants of FAM136A-HA stably and inducibly in a FAM136A KO (**Figure S2**) background (**Figure 4A**). One variant lacked all nine cysteines of FAM136A (9CA), while the two others lacked the four cysteines of one of the twin-CX_3_C motifs of the protein (CA-1, CA-2), respectively. When we tested the cellular localization of these proteins by immunofluorescence, we found that, in steady state, all cysteine mutant variants failed to co-localize with mitochondria. Instead, these variants exhibited weak punctuate cytosolic staining (**Figure 4B**). As explanation for the mislocalization of FAM136A cysteine variants, we found that FAM136A-9CA exhibited an impaired import compared to the WT (**Figure 4C**), indicating that cysteine oxidation in FAM136A by MIA40 is required for efficient import of the protein into mitochondria. Moreover, the levels of the FAM136A variants in intact cells were strongly reduced compared to the WT (**Figure 4D**). This indicates that FAM136A requires all eight conserved cysteines and thus both twin-CX_3_C motifs for IMS import. Lastly, the cysteines of MIA40 substrates are embedded into so-called MISS or ITS motifs [55–57], of which FAM136A also appears to contain one (**Figure 4E**). We generated two MISS/ITS mutant variants (FAM136A M28A/L31A/M32A and M28S/L31S/M32S) and found them indeed to be impaired in mitochondrial import (**Figure 4F**). Thus, FAM136A import depends on its cysteines, a MISS/ITS motif and the activity of the mitochondrial disulphide relay.

**Figure 4.**
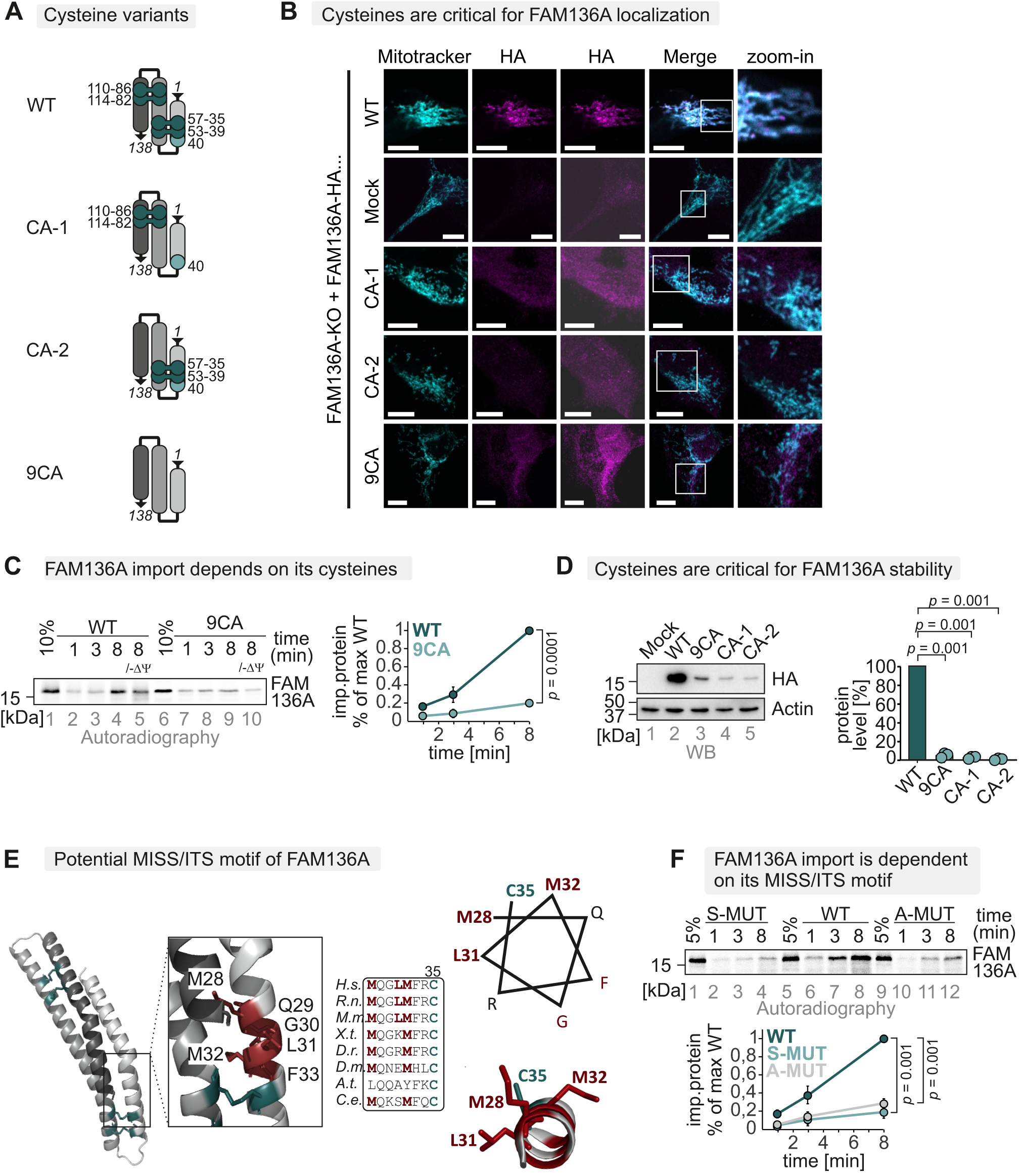
FAM136A cysteines determine its stability. **A.** Scheme of FAM136A cysteine variants. For detailed analysis of cysteines in FAM136A, we generated FAM136A variants: one lacking all cysteines (9CA), one lacking the cysteines 35, 39, 53, 57 (CA-1), and another one lacking the cysteines 82, 86, 110, 114 (CA-2). **B.** Immunofluorescence analysis to localize FAM136A cysteine variants. HEK293 cells stably and inducibly expressing the indicated FAM136A-HA variants were fixated, permeabilized, and stained using a primary antibody against the HA epitope (HA) and mitotracker. Cells were analyzed by fluorescence microscopy. FAM136A-HA WT localizes to mitochondria, while the cysteines variants do not, indicating that cysteines are critical for FAM136A localization. Bar corresponds to 10 µm. **C.** *In organello* import assay with FAM136A cysteine variant to test for its dependence on its cysteines. Experiment was performed as described in Figure 3A. To test for the dependence of FAM136A on its cysteines, a wild-type and a cysteine-to-alanine mutant of FAM136A (9CA, all cysteines mutated) were compared. Import of the cysteine variant was strongly impaired (one-sample t-test). N = 3 replicates **D.** Protein levels in HEK293 cell lines expressing different FAM136A variants. Lysates from different cells were analyzed by reducing SDS-PAGE and immunoblotting. Signals were quantified using ImageLab, and the amount of protein was plotted. All FAM136A variants lacking cysteines were present at very low levels (one-way ANOVA with post hoc Tukey HSD test). N = 3 replicates. **E.** FAM136A contains a patch of hydrophobic amino acids around cysteine 35 that might serve as MISS/ITS facilitating MIA40 interaction. **F.** *In organello* import assay with FAM136A MISS/ITS variant. Experiment was performed as described in Figure 3A. To test for the dependence of FAM136A on its MISS/ITS, a wild-type (WT), a M28A/L31A/M32A and a M28S/L31S/M32S mutant of FAM136A were compared. Import of the MISS/ITS variant was impaired (one-sample t-test). N = 3 replicates

### FAM136A forms a dimer during its biogenesis

When analysing mature FAM136A by gel filtration, we found it to migrate at a higher molecular weight than expected for the monomeric protein (**Figure 5A**, about 30 kDa compared to the expected 15 kDa). Moreover, we could also crosslink FAM136A using the crosslinker disuccinimidylglutarat (DSG), resulting in a specific band at about 35 kDa (**Figure 5B, S3**). In line with this, we could co-precipitate FAM136-HA with FAM136A-FLAG from a cell line stably co-expressing both proteins **(Figure 5C)**. Thus, we propose that mature FAM136A is present in cells as a homo-dimer. This notion is also supported by an Alphafold prediction (**Figure 5D,E**). In this model, aa21-55 of both monomers form the dimerization interface, indicating that dimerization is facilitated by a hydrophobic interaction involving M32, F33 and I54 of both monomers. In addition, the predicted model suggests that R26 may stabilize the dimer through extensive hydrogen bonds with the backbone carbonyls of D42 and A45 of the interacting monomers (**Figure 5F**). These residues are well conserved indicating that FAM136A might be present as a dimer in different species (**Figure 5F**). Since these residues are either part of the MISS signal for FAM136A interaction with MIA40 (**Figure S1**) or very close to one of the disulphide bonds, we assume that dimerization occurs after import and folding of FAM136A.

**Figure 5.**
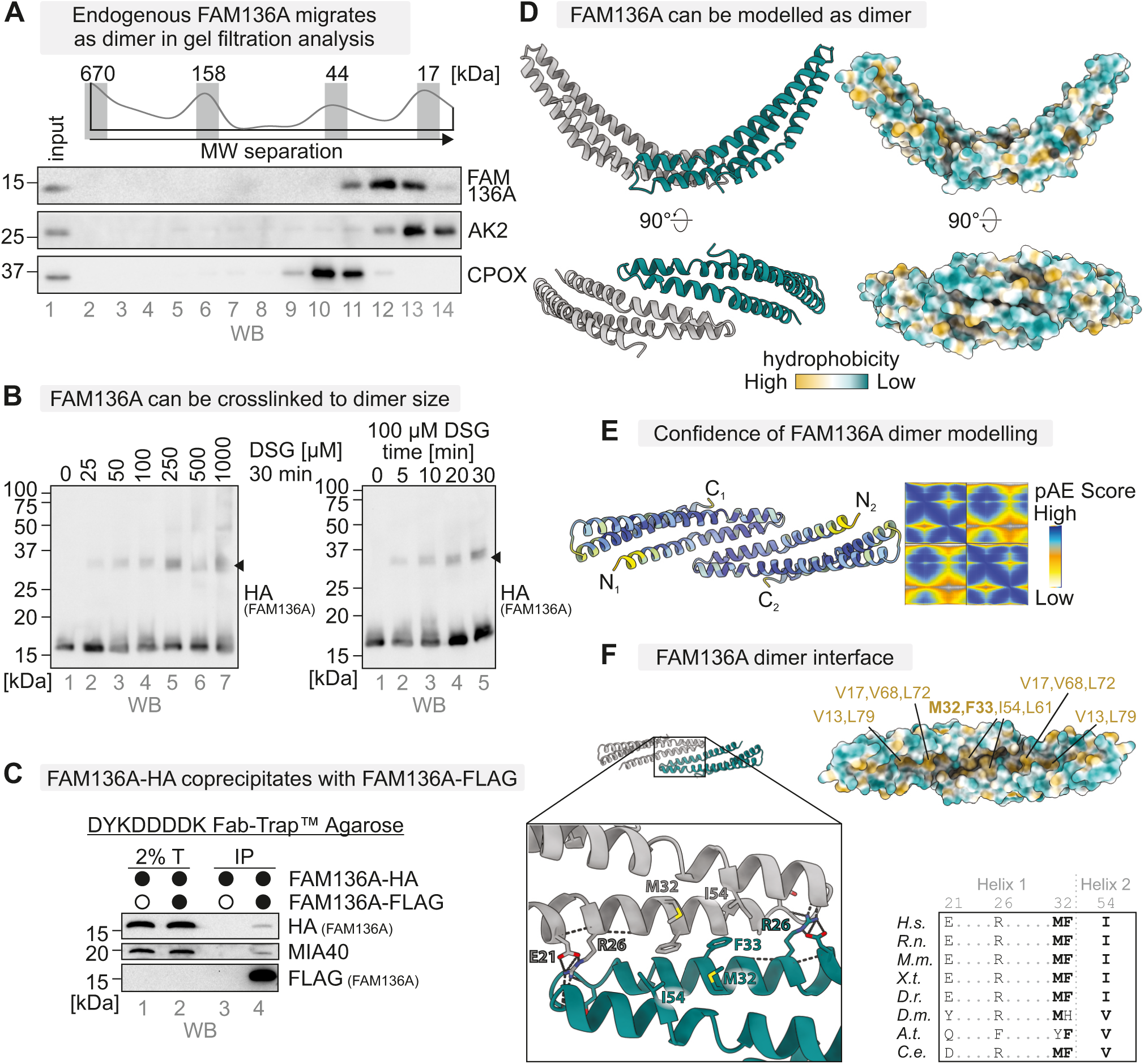
FAM136A forms a dimer during its biogenesis **A.** Gel filtration analysis to assess the native size of endogenous FAM136A in HEK293 cells. HEK293 cells were lysed under native conditions (Triton X-100), and the cleared lysates subjected to gel filtration analysis. Eluted fractions were subjected to TCA precipitation, resuspension in loading buffer containing SDS and DTT, and subsequent immunoblotting against FAM136A, and AK2 and CPOX as controls. As expected, endogenous FAM136A migrates at ca 30 kDa, about twice the expected size (as judged by comparison to protein markers: thyroglobulin, 670 kDa; γ-globulin, 158 kDa; ovalbumin, 44 kDa; myoglobulin, 17 kDa). **B.** Crosslinking analysis of FAM136A-HA. HEK293 cells stably and inducibly expressing FAM136A were incubated for the indicated amounts and times with the crosslinker disuccinimidylglutarat (DSG). The crosslinking reaction was stopped by addition of TRIS. Cells were lysed in loading buffer containing SDS and DTT, and analysed by subsequent immunoblotting against HA. FAM136A can be partially crosslinked to a band of about 30-35 kDa in size (*black arrowhead*). **C.** Analysis of FAM136A homodimerization. HEK293 cells stably and inducibly expressing both, FAM136A-FLAG and FAM136A-HA, were lysed under native conditions and FAM136A-FLAG was precipitated. The lysate was analysed by immunoblotting against HA and FLAG tag. MIA40 served as positive control. The coprecipitation of FAM136A-HA with FAM136A-FLAG together with the sizes determined in (A.) and (B.) indicates the presence of FAM136A as homodimer in intact cells. **D.** Alphafold model of the FAM136A dimer. **E.** AlphaFold model of dimeric FAM136A, showing the prediction of a FAM136A dimer at high confidence. Model colored according to pLDDT score (yellow = low, blue = high). pAE scores of the predicted dimerization of FAM136A. High confidence is assigned to the N-terminal 60 aa taking part in the dimer interface. **F.** Enlarged view of the dimer interface, highlighting the amino acids predicted to stabilize the dimer and conservation of dimer interface. A hydrophobic interface is made up of M32, F33 and I54 of both protomers. R26 forms extensive intra-and intermolecular hydrogen bonds. Key residues shown in ball-and-stick representation. Dashed lines: hydrogen bonds.

### Loss of FAM136A results in loss of selected IMS proteins and increased aggregation of the IMS protein HAX1

Given the critical role of the mitochondrial disulphide relay and many of its substrates for mitochondrial proteostasis [14], we sought to determine the cellular consequences of FAM136A defects. To this end, we acutely transduced cells with lentiviral CRISPR-Cas9 constructs targeting FAM136A and then traced cellular fitness over time in glucose-containing medium [10] (**Figure 6A**). This revealed a notable fitness defect in HEK293 cells depleted of FAM136A (**Figure 6B**), which was even more pronounced in HAP1 cells (**Figure 6C**). In a complementary proliferation experiment using siRNAs (**Figure 6D**), we confirmed the decreased cell proliferation of FAM136A-depleted HEK293 cells (**Figure 6E**). Notably, membrane potential of FAM136A-depleted cells did not substantially change (**Figure S4**).

**Figure 6.**
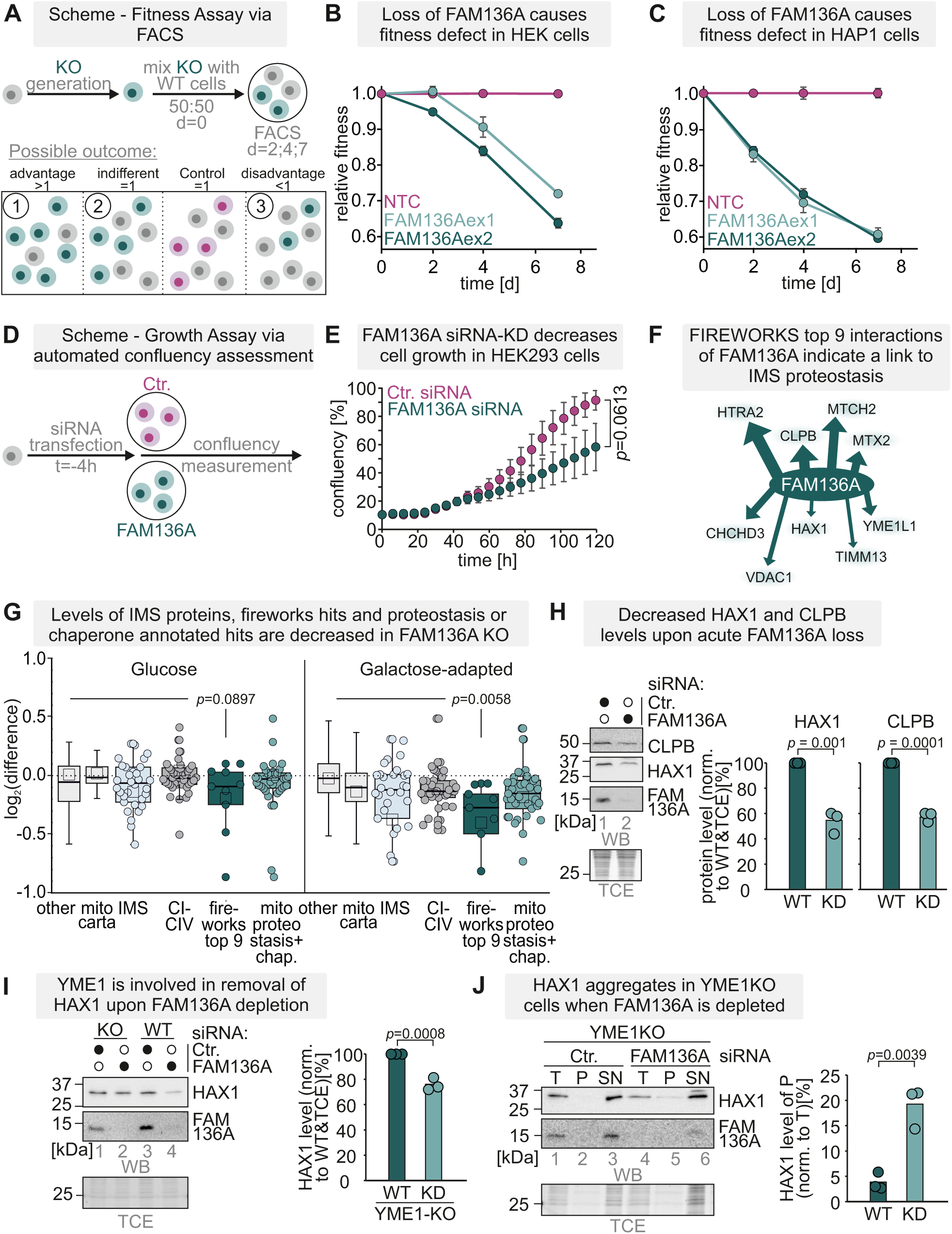
Loss of FAM136A results in diminished cell fitness and IMS proteostasis effects **A.** Layout of competitive fitness experiment. **B,C.** Analysis of competitive cellular fitness upon acute loss of FAM136A in HEK293T **(B)** and HAP1 **(C)** cells. Cells were treated with two different sgRNAs to ablate FAM136A for three days. Subsequently, their fitness was assessed upon growth in glucose-containing medium in a competitive proliferation assay by flow cytometry (see methods). **D.** Layout of proliferation experiment. **E.** Analysis of cell proliferation upon acute loss of FAM136A in HEK293 cells. Cells were treated with siRNA to deplete FAM136A. Subsequently, their proliferation on glucose-containing medium was assessed in a high-throughput microscopy assay by automatically scoring their confluency. N=3 (one-sample t-test) **F.** Coessentiality analysis of FAM136A suggests its functioning in IMS proteostasis networks. **G.** Whole cell proteomes of FAM136 KO cells grown on glucose-or galactose-grown medium. Cells were lysed under native conditions and analyzed by label-free mass spectrometry. N = 4 biological replicates, an unpaired one-sample two-sided Student’s t-test was applied (p<0.01, log2-enrichment >1). **H.** HAX and CLPB levels upon FAM136A loss. Lysates from HEK293 cells depleted for FAM136A by siRNA were analyzed by reducing SDS-PAGE and immunoblotting. Signals were quantified using ImageLab, and the amount of protein was plotted. HAX1 and CLPB levels are reduced in FAM136A siRNA-treated cells (one-way ANOVA with post hoc Tukey HSD test). N = 3 replicates. **I.** HAX1 in YME1L KO cells. Lysates from YME1L KO cells depleted for FAM136A by siRNA were analyzed by reducing SDS-PAGE and immunoblotting. Signals were quantified using ImageLab, and the amount of protein was plotted. HAX1 was reduced in FAM136A siRNA-treated cells and its levels were recovered in YME1L KO cells (one-way ANOVA with post hoc Tukey HSD test). N = 3 replicates. **J.** Protein aggregation assay. Cells were opened and separated into soluble (S) and insoluble (P) components, and analysed by SDS-PAGE and autoradiography. Absence of FAM136A increased the amounts of insoluble HAX1 in YME1L KO cells (one-way ANOVA with post hoc Tukey HSD test). T, total; TSP, total-supernatant-pellet assay. N = 3 replicates.

Coessentiality network analyses using the FIREWORKS and DEPMAP tools puts FAM136A at the center of different proteostasis components acting in the IMS including CLPB, HTRA2, YME1L, HAX1 and MTCH2 (**Figure 6F**) (Amici *et al*, 2021)[58]. In addition, a recent study demonstrated instability of IMS proteins such as HAX1 and ENDOG upon FAM136A depletion in K562 cells [46]. Collectively, this might hint to a role of FAM136A in IMS proteostasis organization. We thus tested whether also in our system FAM136A contributes to IMS proteostasis. Whole cell proteome data from FAM136A KO cells indicated a significant decrease of the top-9 genetic interactors of FAM136A including HTRA2, HAX1, and CLPB (**Figure 6G**). In line, we found the levels of HAX1 and CPLB lowered upon siRNA-mediated depletion of FAM136A (**Figure 6H**). Loss of the IMS protease YME1L recovered HAX1 levels (**Figure 6I**). However, under these conditions, we found in an *in-cell* aggregation assays that absence of FAM136A led to a shift of HAX1 towards the pellet fraction (**Figure 6J**). Collectively, these findings imply that FAM136A counteracts aggregation of specific proteins.

### Loss of FAM136A results in ISR induction and impaired cellular fitness

We had previously observed that impaired proteostasis in the IMS triggers the ISR [11]. To assess whether FAM136A deficiency might engage this signaling axis, we examined the ISR transcription factors ATF4 and CHOP, as well as the ISR-inducible cytokine GDF15 [59]. Indeed, following a lag phase, their expression was induced upon FAM136A loss in both HEK293 and HAP1 cells (**Figure 7A-D**). Interestingly, HAP1 cells displayed both a stronger ISR induction as well as a greater fitness defect following FAM136A depletion, possibly owing to their haploid nature not permitting heterozygous FAM136A mutation. Given the context dependency of ISR effects on cellular fitness [8–12], we tested whether the fitness defect observed upon FAM136A depletion was affected by ISR signaling. While co-deletion of DELE1 or HRI with FAM136A blunted the induction of the tested ISR markers, it did not modulate the fitness phenotype (**Figure S5**). These findings indicate that, although loss of FAM136A robustly activates the mitochondrial ISR, the observed growth defects are likely driven chiefly by a deeper-rooted collapse of mitochondrial proteostasis in the absence of this protein.

**Figure 7.**
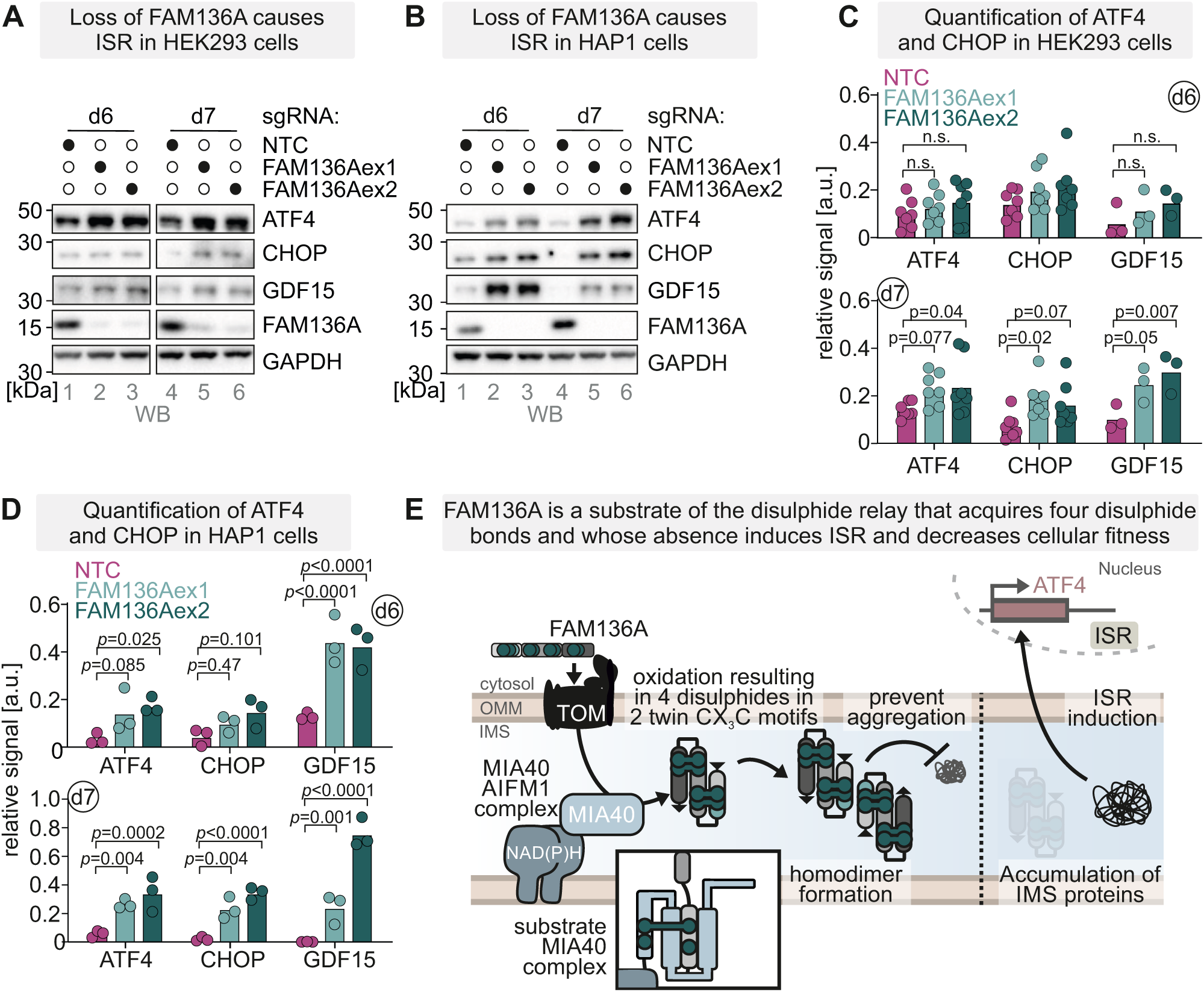
Loss of FAM136A results in ISR induction **A,B.** Analysis of ISR induction upon acute loss of FAM136A in HEK293T **(A)** and HAP1 **(B)** cells. Cells were treated with two different sgRNAs to ablate FAM136A. After 3 days the competitive cell fitness assay was started, and after further 6-and 7-days growth on glucose-containing medium, cells were lysed and analysed by immunoblotting for the indicated proteins. **C.,D.** Quantification of results shown in **(A,B)**. N = 7 replicates for ATF4 and CHOP in HEK293T. N = 3 replicates for all others. Two-way ANOVA. **E.** Model of FAM136A as a disulphide relay substrate involved in IMS proteostasis. *See text for details*

## CONCLUSION

We characterized FAM136A as a new disulphide relay substrate that contributes to IMS proteostasis maintenance (**Figure 7E**). The FAM136A cysteine residues are present in two twin-CX_3_C motifs and are oxidized at steady state. This oxidation is mediated by MIA40 during FAM136A import. They help to retain the protein in the IMS and allow mature FAM136A to homodimerize in cells. The predicted dimerization thereby involves hydrophobic interactions within the segments aa21-55 of each monomer. Absence of MIA40 and AIFM1, the disulphide bond-forming cysteines, or the MISS/ITS motif in FAM136A impairs import and decreases the levels of the protein. Collectively, FAM136A behaves in all parameters like a classical MIA40 substrate:

FAM136A structural motifs indicate resemblance with the family of small TIMM proteins. These proteins have holdase function and contribute to the shuttling of hydrophobic precursors across the IMS including β-barrel proteins to the OMM and mitochondrial carriers to the IMM, respectively. Similarly, FAM136A seems to be important for maintaining proteins like HAX1 in solution and preventing their degradation. In addition to impaired proteostasis, acute loss of FAM136A resulted in decreased fitness in different cellular systems, and triggered an ISR (**Figure 7E, S5J**). A decreased fitness upon FAM136A loss was also reported in two previous studies [46],[47] and is in line with the link of FAM136A to Meniere’s disease [39].

The reported truncated FAM136A patient variant (Q76*) is not imported into the IMS and thus phenocopies the loss of the protein [46]. Notably, the gnomAD database indicates that loss of FAM136A generally is tolerated. This is in line with previous results from *C. elegans*, where depletion of the FAM136A ortholog did not affect lifespan and caused only slight changes in motility and behaviour [60]. Likewise, we observed adaptation effects of FAM136A-KO HEK293 cells; when they were cultured for extended times, they did not exhibit a growth disadvantage compared to WT cells anymore (**Figure S6**).

Altogether, this positions FAM136A as a guardian of mitochondrial proteostasis through a critical effect on a subset of IMS proteins, whose loss acutely provokes pronounced functional deficits in the organelle that can gradually be compensated for under chronic conditions.

## MATERIALS AND METHODS

### Plasmids, cell lines, antibodies and chemical treatments of cells

For plasmids, cell lines, antibodies and other tools used in this study, see **Tables S1-S5**. Cells were cultured in DMEM or IMDM (HAP1) supplemented with 10% fetal calf serum (FCS) at 37°C under 5% CO_2_.

For the cycloheximide (CHX) chase IP experiments, cells were treated for indicated times with 100 µg/ml CHX (dissolved in DMSO). After each time point the cells were washed with ice-cold PBS-NEM prior to harvesting. For the generation of stable, inducible cell lines the HEK293 cell line–based Flp-In T-REx-293 cell line was used with the Flp-In T-REx system (Invitrogen).

### CRISPR-Cas9-based generation of FAM136A HEK293 knockout cell lines

For the generation of FAM136A knockout cells, guide RNA Sequence targeting human FAM136A was cloned into the pSpCas9(BB)-2A-GFP (PX458) vector, which was a gift from Feng Zhang (Addgene plasmid # 48138) {Ran, 2013 #36The guide RNA sequence (Guide:5’-ACCGTCCGGAAGATGCAGGTAGCG-3’) was used. HEK Flp-In T-REx-293 cells were transfected with the plasmid using polyethylenimine (PEI; Thermo Fisher Scientific). After 24 h, GFP-positive cells were collected via FACS and single cell clones were seeded onto 96-well plates (1 cell / well). Clonal cell lines were screened for FAM136A protein expression by western blotting.

### Complementation of FAM136A CRISPR clones

For the complementation of CRISPR clones with different FAM136A variants, the inducible Flp-In T-REx System was used. All FAM136A constructs were cloned into the pcDNA5 FRT-TO vector and co-transfected with the pOG44 Vector into the different CRISPR clones by using the transfection reagent FuGene, according to the manufacturer’s guideline. Positive clones were selected with glucose-containing medium (DMEM supplemented with 1 mM sodium pyruvate, 1 x nonessential amino acids, 10% FCS and 500 mg/ml Pen/Strep, 50 µg/ml Uridine) containing 10 µg/ml Blasticidin and 100 µg/ml Hygromycin.

### Acute FAM136A depletion and competitive fitness assay

HEK293T and HAP1 cells were each infected with a lentiviral construct encoding Cas9, mScarlet, blasticidin resistance, as well as an sgRNA targeting FAM136A exon 1 (5’-GACTCCACCGCCTCCTGCACC-3’), exon 2 (5’-GCTGGTGCACCTGCTTCATGG-3’), or a non-targeting control (NTC) sgRNA (5’-GGTATGTCGGGAACCTCTCC-3’). To generate virus, HEK293T cells were transfected with the respective lentiviral plasmid alongside pAdvantage, pCMVd8.2dVPR and VSV.G. HEK293T and HAP1 cells were transduced following standard procedures, selected with 10 µg/mL (HEK293T) or 18 µg/mL (HAP1) blasticidin for two days and recovered for one day. For fitness measurements, the transduced cells were then mixed with wild-type cells of the respective cell line and seeded in triplicates. The percentage of sgRNA-containing cells (mScarlet^+^) was followed over time by flow cytometry and visualized relative to the day of seeding (d0) and the non-targeting control sgRNA. To measure ISR induction, transduced cells were harvested six-or seven-days post transduction as indicated and analysed by immunoblotting. ISR induction in DELE1/HRI deficient cells was likewise measured by transduction of respective knockout cell lines [8].

For fitness measurements in DELE1/HRI and FAM136A double-KO cells. HEK293T and HAP1 cells were infected either with a lentiviral construct encoding Cas9, GFP, puromycin resistance, as well as an sgRNA targeting DELE1 (5’-GAGCGACATGTGGCGCCTCCC-3’) or HRI (5’-GCCATCGACTTTCCCGCCGA - 3’), or with a construct encoding Cas9, puromycin resistance and a non-targeting control sgRNA (5’-GGTATGTCGGGAACCTCTCC-3’). Virus was generated as described above and transduced HEK293T or HAP1 cells were selected with 2 μg/ml (HEK293T) or 1 μg/ml (HAP1) puromycin for two days. DELE1 or HRI knockout cells were mixed with the non-fluorescent sgNTC cells and transduced with lentiviral sgRNA constructs targeting FAM136A or sgNTC as above. The percentage of sgFAM136A/sgNTC-containing cells (mScarlet^+^) was followed over time by flow cytometry and visualized separately for sgDELE1/sgHRI (GFP^+^) or wildtype cells relative to the day of seeding (d0) and the non-targeting control sgRNA.

### Acute siRNA-mediated FAM136A depletion

SiRNA mediated knockdown of FAM136A was performed by reverse transfection. Therefore 30 pmol RNAi duplex (FAM136A or Ctr.) were diluted in 500 µl opti-MEM I medium without serum in the well of the plate (6-well). 5 µl Lipofectamine RNAiMAX was added to each well, gently mixed and incubated for 10 min at RT. HEK FLP/IN T-Rex cells were dilute in complete medium without antibiotics that 2.5 ml contains 250,000 cells. Cells were growth for 72h to achieve knockdown of FAM136A.

### Cell proliferation assay to test for growth on different carbon sources

For cell proliferation assay recorded with the cytosmart omni 15,000 cells were seeded in 48-well dish and incubated at 37°C. For siRNA experiments, cells were seeded after transfection with siRNA as described above. After 48h, the medium was exchanged with DMEM containing galactose (DMEM supplemented with 4.5 g/l galactose, 1 mM sodium pyruvate, 1 x non-essential amino acids, 10% FCS and 50 µg/ml Uridine). Every 6h the coverage of each well was scanned for 6 days using the cytosmart omni.

### Membrane potential measurement

HEK293T and HAP1 cells were each infected with a lentiviral construct encoding Cas9, GFP, puromycin resistance, as well as an sgRNA targeting FAM136A exon 1 (5’-GACTCCACCGCCTCCTGCACC-3’), exon 2 (5’-GCTGGTGCACCTGCTTCATGG-3’), or a non-targeting control (NTC) sgRNA (5’-GGTATGTCGGGAACCTCTCC-3’). To generate virus, HEK293T cells were transfected with the respective lentiviral plasmid alongside pAdvantage, pCMVd8.2dVPR and VSV.G. HEK293T and HAP1 cells were transduced following standard procedures and selected with 2 µg/mL (HEK293T) or 1 µg/mL (HAP1) puromycin for two days. For membrane potential measurements, the transduced cells were mixed with wild-type cells of the respective cell line and seeded in triplicates.

On days 6 and 7 post transduction, the cells were treated for 30 min with 100 μM TMRM in the respective medium, or with 100 μM TMRM and 20 μM CCCP as a positive control for mitochondrial depolarization. TMRM fluorescence was measured by flow cytometry and normalized to the non-transduced control cells from the same well.

### Image acquisition

#### Microscope image acquisition

For the image acquisition the microscope LSM 980 with Airyscan 2 and multiplex from Carl Zeiss Microscopy was used with a Plan-Apochromat 63x/1,4 Oil DIC objective and the GaAsP-PMT, Multi-Alkali-PMT detector. The cells were imaged at room temperature with oil as imaging medium. The following fluorochromes were used: Mitotracker CMXRos and Alexa Fluor488.

Images were displayed using the acquisition software ZEN 3.3. and were processed using the software OMERO.insight.

#### Western Blot image acquisition

The immunoblotting images were detected using the ChemiDoc Touch Imaging system (Bio-Rad). The autoradiography images were detected using the image analyzer Typhoon FLA 7000.

#### Native and denaturing immunoprecipitation (IP)

To detect protein-protein interactions, native or denaturing IP were performed. Corresponding cell lines were seeded on a 10 cm dish and were grown until confluency of 90%. Before IP was performed, inducible cell lines were induced with doxycycline overnight or for three days. Then the cells were washed with 5 ml ice-cold PBS containing 20 mM NEM and were incubated with another 5 ml ice-cold PBS containing 20 mM NEM for 15 min on ice. The cells were scratched off and pelleted at 700 x g for 3 min at 4°C. For native IP, the pellet was resuspended in 1000 µl native IP buffer (100 mM Na-P_i_ pH 8.1, 100 mM NaCl, 1% Triton X-100) and was incubated for at least 30 min on ice. For denaturing IP, the pellet was resuspended in 200 µl denaturing IP buffer (30 mM Tris pH 8.1, 100 mM NaCl, 5mM EDTA) and 50 µl 10% SDS was added. The denaturing IP samples were sonified at maximum amplitude. After cooling down, 750 µl Denaturing buffer +2.5 % Triton X 100 was added and incubated for at least 30 min on ice. After that, the samples were centrifuged at 21,817 x g for 1 h at 4°C. The supernatant was collected and incubated with 10 µl equilibrated HA beads (monoclonal anti-HA-Agarose produced in mouse) or with Streptactin beads overnight at 4 °C. Then, the beads were washed 4 times with 1 ml washing buffer containing triton X-100 (100 mM Na-P_i_ pH 8.1, 1% Triton X-100, 100 mM NaCl for native IP; 30 mM Tris pH 8.1, 100 mM NaCl, 5 mM EDTA, 1% Triton X-100 for denaturing IP) and 1 x with washing buffer without Triton X-100 with a centrifugation step at 2,000 x g for 1 min at 4°C. After the last centrifugation step, the beads were dried completely and 20 µl reducing Laemmli was added and the samples were boiled for 10 min. The samples were analysed by SDS-PAGE and Western Blot.

#### Isolation of crude mitochondria from HEK293 cells

Isolation of crude mitochondria from HEK293 cells was performed as described in {Murschall, 2021 #49}. In short, cells were cultivated for 4 days on 15 cm dishes. For harvesting, the cells were washed 2 times with 10 ml ice-cold 1 x PBS and scraped off using a cell scraper. Afterwards the cells were centrifuged at 500 x g for 5 min at 4°C. The pellets were resuspended in 5 ml 1x M buffer (220 mM mannitol, 70 mM Sucrose 5 mM HEPES-KOH, pH 7.4 1 mM EGTA-KOH, pH 7.4) containing 1 x Complete TM Protease Inhibitor Cocktail. The cells were homogenized using a precooled potter homogenizer (1,000 rpm, 15 strokes). The homogenized cells were pelleted at 600 x g for 5 min at 4°C. The supernatant containing the crude mitochondria was centrifuged at 8,000 x g for 10 min at 4°C. The pellet was washed with 2 ml ice-cold 1x M buffer (without the complete protease inhibitor cocktail). The crude mitochondria were pelleted at 6,000 x g for 10 min at 4°C and the supernatant was carefully removed. The pellet was resuspended in 400 µl and the concentration was measured using the BCA Reagent ROTI® Quant Assay according to the manufactureŕs instructions.

#### Submitochondrial fractionation of isolated mitochondria

For subcellular fraction of FAM136A 40 µg crude mitochondria was centrifuged at 10,000 x g for 5 min at 4°C. Each pellet was resuspended in 95 µl of corresponding buffer (<u>isotonic</u> buffer: 1 x M buffer (220 mM mannitol 70 mM Sucrose 5 mM HEPES-KOH, pH 7.4 1 mM EGTA-KOH, pH 7.4); <u>hypotonic</u> buffer: 10 mM HEPES pH 7.4; <u>TX-100</u> buffer: 1x M buffer,1% TX-100; each buffer containing either 40µg/ml PK or no PK) by pipetting up and down using a 200 µL pipet tip that was cut. The samples were incubated for 5 min at 21 °C and after the incubation time 2,5 µl of 0.2 M PMSF (final concentration 5 mM) was added to all 6 samples and incubation was allowed for a further 5 min on ice. Then, the samples were centrifuged at 10,000 x g for 5 min and the pellets were resuspended with 100 µl of each buffer (isotonic or hypotonic) containing 1 mM PMSF. 4x Laemmli buffer and DTT was added to a final concentration of 50 mM DTT. All samples were boiled at 95°C for 5 min. The samples were analysed by SDS-PAGE and Western Blot.

#### Protein import assay into isolated mitochondria

The import assay was performed as described in [54]. After isolation, 20 µg of mitochondria per import reaction were centrifuged at 8,000 x g for 5 min at 4°C. The resulting pellet was carefully mixed and resuspended with 4 µl [^35^S]-labelled precursor lysate (prepared according to manufactureŕs protocol and [54]) using a cut 10 µl pipette tip. For the CCCP control, 1 µl of 20 mM CCCP solution was added and incubated on ice prior to import reaction. The import reactions were incubated at 30°C while shaking at 600 rpm and were stopped at different time points by putting on ice. The samples were directly centrifuged at 8,000 x g 5 min at 4°C. Then, the samples were treated with proteinase K (PK) by resuspending mitochondria in 400 µl of 1 x M buffer (220 mM mannitol 70 mM Sucrose 5 mM HEPES-KOH, pH 7.4 1 mM EGTA-KOH, pH 7.4) containing 20 µg/ml PK. The samples were incubated for 20 min on ice and the digestion was stopped by adding 2 µl of 200 mM PMSF (1 mM final).

After another centrifugation step at 10,000 x g for 5 min at 4°C the pellets were washed with 400 µl buffer M containing 1 mM PMSF. The resulting pellets were resuspended in 30 µl reducing Laemmli buffer. The samples were boiled at 96°C for 5 min. In case the samples turned yellow, the samples were pH adjusted with a 1 M Tris solution (unbuffered). The samples were analysed by SDS-PAGE and autoradiography.

#### Determination of protein interaction partners by proteomics (interactome analysis)

For the interactome data an in-gel digest was performed. The native IP was performed as described in the respective section. After the native IP was performed the beads were dried and boiled in 20 µl reducing Laemmli buffer (without bromphenolblue) for 10 min. The samples were reduced by addition of DTT with a final concentration of 5 mM and incubated at 56 °C for 30 min. Free cysteine thiols were alkylated by addition of CAA to a final concentration of 40 mM to the samples which were then incubated for 30 min at room temperature in the dark. The samples were run on SDS-PAGE until the samples migrated for 1 cm into the separation gel. Then the gels were fixed for 1 h in fixing solution (10% Acetic acid / 20% Methanol in water). The gel bands were cut in smaller pieces and were transferred to individual tubes. 100 µl of 50 mM ABC/50% Acetonitrile was added to the gel pieces and were incubated for 20 min at room temperature. The solution was exchanged with fresh 50 mM ABC/50% Acetonitrile and remaining solution was discarded after 20 min incubation. The gel pieces were covered with 100 µl acetonitrile and incubated for 10 min. The gel pieces were then dried in a speedvac for approximately 5 min. A digestion solution of 10 ng/μL of 90 % trypsin and 10 % LysC in 50 mM ammonium bicarbonate (ABC) was added to the gel pieces until the gel pieces were fully covered. The gel pieces were incubated for 30 min at 4°C with the digest solution.

After the incubation time, excessive digest solution was removed and 50 mM ABC buffer was used to cover the gel pieces. The samples were incubated overnight at 37°C while shaking at 750 rpm. The next day, the supernatant of the gel pieces were transferred into new tubes. The gel pieces were covered with 100 µl 30% ACN/3% TFA and incubated for 20 min at room temperature. The extract was combined with the supernatant of the previous step. The gel pieces were covered with 100 µl 100% acetonitrile and again incubated for 20 min at room temperature. The extract was also combined with the supernatant from the previous step and the organic solvents of the samples were reduced in the speedvac until a remaining volume of 50 µl was reached. The samples were acidified by addition of formic acid to a final concentration of 1% and the STAGE tip purification protocol as it is described in the section Samples were analyzed by the CECAD Proteomics Facility on a QExactive Plus (QEP) mass spectrometer coupled to an Easy nLC 1000 or on an Orbitrap Exploris 480 mass spectrometer coupled to an Vanquish neo in trap-and-elute setup (all Thermo Scientific). Samples were either directly loaded onto an in-house packed analytical column (30 cm length, 75 µm inner diameter, filled with 2.7 µm Poroshell EC120 C18, Agilent, Easy nLC setup) or first onto a precolumn (Acclaim 5µm PepMap 300 µ Cartridge) with a flow of 60 µl/min before reverse-flushed onto an identical main column. Peptides were chromatographically separated on a linear gradient between 3% and 30% in 45 min followed by column washing and reequilibration. The mass spectrometers were operated in data-dependent acquisition in Top10 mode 8(QEP) or with a cycle time of 1 s (Exploris) with MS1 scans acquired from 350 m/z to 1400 m/z at 70k (QEP) or 60k (Exploris) resolution and an AGC target of 300%. MS2 scans were acquired at 17.5k (QEP) or 15 k (Exploris) resolution with a maximum injection time of 118 ms and a normalized AGC target of 50% in a 2 Th window and a fixed first mass of 110 m/z. All MS1 scans were stored as profile, all MS2 scans as centroid.

All mass spectrometric raw data were processed with MaxQuant (version 2.4, Tyanova et al. 2016a) using default parameters against a Swissprot Human canonical database (UP5640, downloaded 04/01/2023) with the match-between-runs option enabled between replicates. Follow-up analysis was done in Perseus 1.6.15 (Tyanova et al. NatMet). Protein groups were filtered for potential contaminants and insecure identifications. Remaining IDs were filtered for data completeness in at least one group and missing values imputed by sigma downshift (0.3 σ width, 1.8 σ downshift). Afterwards, FDR-controlled two-sided t-tests were performed.

#### Determination of cellular protein levels by quantitative label-free proteomics

For quantitative label-free proteomics, the experiments were performed as described in [48]. Respective cells were seeded on a 6 well dish. The next day the cells were washed with PBS and harvested by cell scraper and centrifugation at 500 x g for 5 min. The pellets were resuspended in 20 µl lysis buffer (4% SDS in PBS containing protease inhibitor) and were sonified. Afterwards the samples were boiled for 5 min at 96°C. After cooling down, 80 µl ice-cold acetone was added and stored at-80°C overnight. The next day, TCA precipitations were thawed, and the samples were centrifuged for 15 min at 16.000 x g. The resulting pellets were washed with 500 µl acetone and then air-dried. The pellets were then resuspended in 50 µl 8 M urea in TEAB buffer supplemented with protease inhibitor cocktail and were sonified. The samples were centrifuged for 15 min at 20,000 x g. The supernatant was transferred into a new tube and the protein concentration of the samples were determined by Pierce Protein Assay Reagent. The assay was performed according to the manufactureŕs protocol and the concentration was measured at 600 nm. 50 µg of each sample was transferred to a new reaction tube and filled up to 40 µl with the Urea/TEAB buffer. Then, DTT with a final concentration of 5 mM was added and the samples were incubated for 1 h at 37°C. Next, chloroacetamide (CAA) with a final concentration of 40 mM was added to the samples and were incubated for 30 min in the dark. For the digest of the peptides, first Lysyl Endopeptidase with an enzyme to substrate ratio of 1:75 was used and the samples were incubated for 4 h at 25°C. For the trypsin digest the samples were first diluted with TEAB buffer to reach a urea concentration below 2 M and then trypsin with an enzyme to substrate ratio of 1:75 was added. The samples were incubated at 25 °C overnight. In the last step, the samples were purified by STAGE tips which were equilibrated with methanol and buffers containing 0.1 % formic acid and 80 % acetonitrile. The samples were loaded on the STAGE tips and were washed with buffers containing 0.1 % formic acid and 80 % acetonitrile. The STAGE tips were completely dried and until measurement stored at 4 °C. Samples were analyzed by the CECAD Proteomics Facility on an Orbitrap Exploris 480 mass spectrometer equipped with a FAIMSpro differential ion mobility device that was coupled to an UltiMate 3000 (all Thermo Scientific). Samples were loaded onto a precolumn (Acclaim 5µm PepMap 300 µ Cartridge) for 2 min at 15 ul flow before reverse-flushed onto an in-house packed analytical column (30 cm length, 75 µm inner diameter, filled with 2.7 µm Poroshell EC120 C18, Agilent).

Peptides were chromatographically separated at a constant flow rate of 300 nL/min and the following gradient: initial 6% B (0.1% formic acid in 80 % acetonitrile), up to 32% B in 72 min, up to 55% B within 7.0 min and up to 95% solvent B within 2.0 min, followed by column wash with 95% solvent B and reequilibration to initial condition.

The FAIMS pro was operated at-50V compensation voltage and electrode temperatures of 99.5 °C for the inner and 85 °C for the outer electrode.

MS1 scans were acquired from 399 m/z to 1001 m/z at 15k resolution. Maximum injection time was set to 22 ms and the AGC target to 100%. MS2 scans ranged from 400 m/z to 1000 m/z and were acquired at 15 k resolution with a maximum injection time of 22 ms and an AGC target of 100%. DIA scans covering the precursor range from 400 - 1000 m/z and were acquired in 60 x 10 m/z windows with an overlap of 1 m/z. All scans were stored as centroid.

Samples were analyzed in DIA-NN 1.8.1 (Demichev 2020). A Swissprot Human canonical database (UP5640, downloaded 04/01/2022) was used for library building with settings matching acquisition parameters and the match-between-runs function enabled. Here, samples are directly used to refine the library for a second search of the sample data. DIA-NN was run with the additional command line prompts “— report-lib-info” and “—relaxed-prot-inf”. Further output settings were: filtered at 0.01 FDR, N-terminal methionine excision enabled, maximum number of missed cleavages set to 1, min peptide length set to 7, max peptide length set to 30, min precursor m/z set to 400, max precursor m/z set to 1000, cysteine carbamidomethylation enabled as a fixed modification. Afterwards, DIA-NN output was further filtered on library q-value and global q-value <= 0.01 and at least two unique peptides per protein using R (4.1.3). Finally, LFQ values calculated using the DIA-NN R-package. Afterwards, analysis of results was performed in Perseus 1.6.15 (Tyanova 2016b). Here, results were filtered for data completeness in corresponding replicate groups and imputed using the minDet function followed by FDR-controlled T-tests.

#### Assay to detect redox states of protein thiols

To assess the redox state of FAM136A, 60,000 cells were seeded on a 24-well dish. The cells were cultivated for two days. Before the cells were harvested, they were treated for 19 h with doxycycline (1 µg/ml) to induce FAM136A expression. The assay was coupled to a cycloheximide (CHX) treatment. Therefore, the cells were treated with 100 µg/ml CHX before they were harvested and modified. For harvesting the cells were washed with 500 µl ice-cold PB. The PBS was removed and 1000 µl 8% ice Cold TCA was added to each well. The cells were scratched off and transferred to 1.5 ml eppis and were stored at-80°C until the liquid was completely frozen. Samples were thawed at RT and pelleted for 15 Min, 13.000xg 4°C. The Supernatant was removed and 900 µl of ice-cold 5% TCA were added. After vortexing the samples were centrifuged again for 15 min, 13.000xg, 4°C. The TCA was removed completely. Non-reducing Laemmli buffer (2% SDS, 60 mM Tris, pH 6.8, 10% glycerol, 0.0025% bromphenol blue) containing 15 mM mmPEG12 was added to obtain “steady state” samples. The “maximum reduced” and “unmodified” samples contain in addition to Laemmli buffer 10 mM Tris(2-carboxyethyl) phosphine (TCEP) and the alkylating reagent for the “maximum shift” and no reagent for the “unmodified” sample. The maximum reduced samples were boiled for 15 min at 96°C. All samples were sonicated and were analysed by SDS-PAGE and Western Blot.

#### Gel filtration

Analytical size-exclusion chromatography was performed under native conditions to examine the oligomeric state of the endogenous proteins. Cells were washed with 1x PBS, mechanically detached by scraping and sedimented at 500 x g for 5 min. Cell pellets were resuspended in 1 ml native lysis buffer (100 mM sodium phosphate pH 8.0, 100 mM sodium chloride, 1% (v/v) Triton X-100), supplemented with 200 µM PMSF and incubated for 1 h on ice. The lysate was cleared by centrifugation (20 000 x g, 1 h) and loaded on a HiLoad™ 16/600 Superdex 200 preparation grade gel filtration column installed in a liquid chromatography system (Aekta Purifier) from GE Healthcare. A protein size standard was used as a reference, covering a range from 1.35 kDa to 670 kDa (#1511901, Bio-Rad).

#### Crosslinking of FAM136A with disuccinimidyl glutarate (DSG)

In order to check whether FAM136A can be crosslinked in intact cells, HEK cells were seeded in a 12-well plate until they reach 90 % confluency. Cells were placed on ice and washed twice with ice-cold PBS. Afterwards the DSG-PBS solution with different concentrations of DSG (0, 25, 50, 100, 250, 500, 1000 µM) was added carefully. Incubation occurred at room temperature for 30 min (or was stopped after different time points). The reaction was stopped by removing the crosslinker and washing the cells with 20 mM Tris pH 7.5 and incubation of 20 mM Tris pH 7.5 for 15 min at room temperature. Cells were scratched off and sedimented at 500 x g for 5 min at 4 °C and resuspended in 20 µl 1x reducing SDS sample buffer. The samples were boiled for 10 min at 96 °C and analysed by Western Blot.

#### Preparation of Immunofluorescence (IF) samples

To establish the localization of different FAM136A cysteine mutants, cells were seeded onto poly-L-lysine coated coverslips in 12-well dishes in complete medium. Protein expression was induced by adding 0.1 µg/ml doxycycline for 72 h prior to preparation of IF samples. For Mitotracker staining media was replaced by 1 ml pure media (w/o FCS, P/S) substituted with Mitotracker. Incubation occurred for 1 h at 37°C. Afterwards the media was replaced by complete media followed by an incubation for 30 min at 37 °C. Cells were washed with prewarmed PBS and 1 ml fixation buffer (4 % PFA in PBS) was added and incubated for 15 min @ RT. Cells were washed 3x with PBS and 1 ml fresh blocking buffer (10mM Hepes pH 7.4, 3% BSA, 0.3 % Triton X-100) was added for 1 h at RT. Cells were washed 3x with PBS. 40 µl per coverslip primary antibody solution in blocking buffer (aHA (rabbit) 1:400) was placed onto a parafilm and coverslips were added upside down to the solution for 1 h at RT. Cells were washed 3x with PBS. 40 µl per coverslip secondary antibody solution in blocking buffer (Alexa488 rabbit 1:400) was added onto a parafilm and coverslips were placed upside down to the solution for 1 h at RT in the dark. Cells were washed 3x with PBS and 20 µl prewarmed mixture of Mowiol and DABCO (50 °C) was added onto glass slide and coverslips were placed on top. The samples were dried at 4 °C overnight and sealed with nail polish.

#### In vitro MIA40-GST pulldown

To access the interaction of FAM136A with different MIA40 variants *in vitro,* MIA40-GST WT, C4/53/55S and F68E were expressed and purified and incubated with radiolabelled lysates of FAM136A. MIA40 variants (or pGEX-GP-1 empty vector) were transformed into Rosetta *E. coli* and grown in LB medium with 100 µg/ml ampicillin + 30 µg/ml Chloramphenicol. Protein expression was induced with 0.1 mM IPTG for 5 h at 30 °C and 160 rpm. The bacterial pellets were thawed on ice and resuspended in 15 ml GST buffer (20 mM Tris pH 7.4, 200 mM NaCl + 0.2 mM PMSF). Cell lysis was performed with French press three cycles with 9000 psi. The lysate was centrifuged at 35.000 x g for 20 min at 4 °C. In the meantime, 20 µl glutathione beads per condition were wash together with 1 ml GST buffer (4x) (2 min, 2000g, 4 °C). 1 ml of the supernatant of the different purified variants was added to 20 µl glutathione beads and incubated for overnight at 4 °C. The beads were wash twice with GST-buffer (2 min, 2000g, 4 °C) followed by two washing steps with GST low salt buffer (10 mM NaCl, 20 mM Tris pH=7.4). A quick CBB staining of MIA40 purification was done to see if the different MIA40 variants show equal levels. 6 µl radiolabelled FAM136A-HA lysate was added to the beads and 500 µl IP buffer (100 mM NaCl, 100 mM NaP_i_, 1 % Triton X100) was added. Incubation occurred for 20 min at room temperature followed by a centrifugation step at 1000 x g for 2min at room temperature. The samples were washed 6 x with GST (medium salt) buffer 100 mM NaCl + 20 mM Tris +1 % Triton and 100 µl 1 x Läemmli buffer with 50 mM DTT was added to the beads and incubated for 5 min at 96 °C.

#### Purification of MIA40(C4S)-His

Recombinant proteins were expressed from pET21b-MIA40(C4S) in Rosetta2 E. coli strains. Bacterial growth was conducted in LB media shaking at 37°C and 180 rpm. The expression was induced with 0.5 mM IPTG and incubated for further 3 h. Cells were harvested on ice in PBS and stored at-20°C. The 6xHis-tagged construct was purified by Immobilized Metal Affinity Chromatography using Ni Sepharose (6 FastFlow, GE) in binding buffer (200 mM sodium chloride, 20 mM Tris/Cl pH = 7.4). The bacterial lysate was bound to beads in binding buffer supplemented with 10 mM imidazole at 4°C. Beads were washed with binding buffer supplemented with 20 mM imidazole prior to elution with 150 mM imidazole. Imidazole was removed using PD-10 columns (Cytiva) and the proteins stored at 4 °C.

#### In vitro oxidation kinetics

In vitro oxidation kinetics were carried out at 30 °C and under 1.0 % oxygen. FAM136A was expressed by the TNT Quick Coupled Transcription/Translation System (Promega) according to the manufacturer’s instructions. It was mixed with either buffer (10 mM EDTA, 200 mM NaCl, 20 mM Tris/Cl pH = 7.4) or 30 µM recombinant MIA40(C4S) in buffer. The oxidation was stopped directly by cysteine modification in Laemmli buffer containing 5 mM mmPEG12. As a maximum size shift control, FAM136A was additionally reduced with 5 mM TCEP. All samples were analyzed by non-reducing SDS-PAGE and autoradiography.

#### Fractionation

Cellular fractionation experiments were performed to distinguish between mitochondrial and cytosolic proteins. For this, cells were grown on 15 cm dishes until they reached 80 % confluency. Cells were washed with 10 ml ice-cold PBS and transferred to falcon tubes for centrifugation at 500 x g for 5 min at 4 °C. The cell pellets were resuspended in 2 ml fractionation buffer (20 mM HEPES pH 7.4, 220 mM Mannitol, 70 mM Sucrose, 1 mM EDTA) and homogenized using the potter with 15 strokes at 1100 rpm. Intact cells and cell debris were pelleted at 800 x g for 5 min at 4 °C. The supernatant was transferred into a 2 ml reaction tube and a total sample (T) was taken. The homogenized sample was centrifuged at 13000 x g for 15 min at 4 °C to divide a supernatant (cytosolic) and pellet (mitochondrial). The cytosolic fraction was centrifuged at full speed for 10 min and the supernatant was used for TCA precipitation. The mitochondrial pellet was resuspended in 1 ml fractionation buffer and centrifuged at 13000 x g for 15 min at 4 °C. This washing step was repeated three times before the pellet was resuspended in 80 µl 100 mM Tris/AC pH 8.0. All tree samples (T, cyto, mito) were used for TCA precipitation. Finally, the TCA pellets were resuspended in buffer A and boiled at 96 °C for 5 min.

#### In-cell aggregation assay

SiRNA-mediated knockdown of FAM136A was performed 72 h prior to solubility assay in YME1-KO HEK293 cells. Cells were washed 1x with cold PBS and transferred into a 1.5 ml reaction tube. After centrifugation at 500 g for 5 min at 4 °C, the pellets were resuspended in 75 µl lysis buffer (1 % (v/v) Triton X-100, 200 mM KCl, 30 mM Tris/HCl (pH 7.4), 0.5 mM PMSF, 1x protease inhibitor). 25 µl were used as a “total” (T) sample whereas the remaining 50 µl were incubated for 10 min at 4 °C with occasional vortexing. Cell lysates were centrifuged at 20,000 g for 20 min at 4°C. The supernatant was transferred into a fresh tube and the pellet was washed in 300 µl lysis buffer repeated by additional centrifugation at 20,000 g for 10 min at 4°C. The pellet was finally resuspended in 50 µl of 1 x Solubilisation buffer (5 % SDS, 50 mM triethylammonium bicarbonate (TEAB)) and sonified until the pellet was completely resuspended. Afterwards 18 µl 4 x Laemmli and 3.4 µl 1 M DTT was added and samples were boiled before analysis by SDS-PAGE and Western Blot.

#### Sodium carbonate extraction

In order to access the solubility of mitochondrial proteins, 200 µg freshly isolated mitochondria were centrifuged at 10,000 g for 5 min and resuspended in 500 µl of ice-cold 0.1 M Na_2_CO_3_ solution pH 11.05. Incubation for 1.5 h at 4 °C was performed to open the membranes. A “total (T)” sample was removed. The samples were filled up to 14 ml total volume with Na_2_CO_3_ for ultracentrifugation at 90 000 g (35 000 rpm) for 30 min, 4 °C. The supernatant (14 ml) was collected to perform a TCA precipitation. Both samples (P and SN) were finally resuspended in 131.25 µl MQ (+ 43.7 µl 4xL + 8.75 µl 1 M DTT) and boiled at 96 °C for 5 min.

#### Structure modelling using Alphafold-Multimer

To model the structure of the FAM136A monomer and dimer AlphaFold 3 was used [61]. Predicted best model structures were analyzed and visualized with PyMol or ChimeraX [62].

#### Quantification and statistical analysis

Intensity of autoradiography and immunoblot signals were quantified using ImageQuant and Image Lab (Biorad), respectively. Microscope images were processed with OMERO. Error bars in figures represent standard deviation. The number of experiments is reported in the figure legend. To statistically analyse the import efficiency of two distinct conditions after 30 min of import, a one-sample t-test was performed for each condition. To statistically analyse the FAM136A western blot signal between three different cell lines (WT, KO and complementation), a one-way ANOVA was performed. In cases of significant differences between the cell lines (P < 0.05), a post hoc Tukey HSD test was calculated.

## DATA AVAILABILITY

The datasets generated during and/or analysed during the current study are available from the corresponding author on reasonable request. Data are available via PRIDE database with identifier PXD060952 and PXD060963.

## ACKNOWLEDGEMENTS

The Deutsche Forschungsgemeinschaft (DFG, German Research Foundation) funds research in the Laboratory of JR through the grants RI2150/5 project number 435235019, RTG2550/2 project number 411422114, SPP2453 project number 541742459, CRC1218 - project number 269925409, and CRC1678 – project number 520471345. SP is funded by CMMC core funding (JRG XI), by the DFG - SFB1430 - Project-ID 424228829 and the CANTAR network funded by the Ministry of Culture and Science of the state of Northrhine-Westphalia. Research in the laboratory of LTJ is supported by the European Research Council (ERC-StG 804281 – *SOLID*), the DFG (JA 2873/2-1, project number 470553481 and SPP2453, project number 541592768), the Alfried Krupp von Bohlen und Halbach Foundation, and the Vallee Foundation. HF received additional support by the Boehringer Ingelheim Fonds. We thank Anja Wittmann and Anika Seiler for technical support throughout the project, and Kathrin Ulrich and Matthias Weith for critical reading of the manuscript. We thank the CECAD Proteome and Imaging Facilities for the provision of instrumentation, training, and technical support. This work was supported by the large invest grants INST 1856/71-1 FUGG and INST 216/1163-1 FUGG by the Deutsche Forschungs Gemeinschaft.

## AUTHOR CONTRIBUTIONS

JR and CZ designed the study and planned experiments. CZ designed, cloned the constructs and generated cell lines. CZ performed bioinformatical analyses. CZ and CECAD proteomics facility carried out the mass spectrometry-based proteomics and performed the bioinformatical analysis of the mass spectrometry data. CZ carried out the biochemical experiments. CZ carried out the fluorescence microscopy experiments and the image analysis. RAR performed the *in vitro* oxidation experiment. HF and LTJ conceived, performed, and analysed proliferation and ISR induction experiments upon acute FAM136A loss. SP performed the Alphafold analysis of monomeric and dimeric FAM136A. JR, CZ analysed the data. JR wrote the manuscript with the help and input of all authors.

