## Supplemental Information for "The mitochondrial disulphide relay substrate FAM136A safeguards IMS proteostasis and cellular fitness"

#### TABLE OF CONTENT

##### SUPPLEMENTAL TABLES


Table S1. Cell lines page 3

Table S2. Primers and Plasmids page 4

Table S3. Antibodies page 5

Table S4. Oligonucleotides and siRNAs page 6

Table S5. Further Tools and Equipment page 7


##### SUPPLEMENTAL FIGURES


Figure S1 page 8

Figure S2 page 9

Figure S3 page 10

Figure S4 page 11

Figure S5 page 12

Figure S6 page 14

#### SUPPLEMENTAL TABLES

#### Table S1. Cell lines

| Cell line | Plasmid | Gene | Tag (C-terminal) | Reference |
|---|---|---|---|---|
| **Flp-In T-REx-293** | -- | -- | -- | ThermoFisher R78007 |
| **Flp-In T-REx-293-Mock** | pcDNA5/FRT/TO | -- | -- | (Murschall et al., 2020) |
| **Flp-In T-REx-293-FAM136A knockout** | pSpCas9(BB)2A-GFP | Guide: TCCGGAAGATGCAGGTAGCGGGG | -- | this study |
| **Flp-In T-REx-293- FAM136A knockout-Mock** | pcDNA5/FRT/TO | -- | -- | this study |
| **Flp-In T-REx-293- FAM136A knockout- FAM136A** | pcDNA5/FRT/TO | ORF *FAM136A* | HA | this study |
| **Flp-In T-REx-293- FAM136A knockout- FAM136A -9CA** | pcDNA5/FRT/TO | ORF *FAM136A* ($T_{103}$▶G, $G_{104}$▶C, $T_{115}$▶G, $G_{116}$▶C, $T_{118}$▶G, $G_{119}$▶C, $T_{157}$▶G, $G_{158}$▶C, $T_{169}$▶G, $G_{170}$▶C, $T_{244}$▶G, $G_{245}$▶C, $T_{256}$▶G, $G_{257}$▶C, $T_{328}$▶G, $G_{329}$▶C, $T_{340}$▶G, $G_{341}$▶C) | HA | this study |
| **Flp-In T-REx-293- FAM136A knockout- FAM136A -CA-1** | pcDNA5/FRT/TO | ORF *FAM136A* ($T_{103}$▶G, $G_{104}$▶C, $T_{115}$▶G, $G_{116}$▶C, $T_{157}$▶G, $G_{158}$▶C, $T_{169}$▶G, $G_{170}$▶C) | HA | this study |
| **Flp-In T-REx-293- FAM136A knockout- FAM136A -CA-2** | pcDNA5/FRT/TO | ORF *FAM136A* ($T_{244}$▶G, $G_{245}$▶C, $T_{256}$▶G, $G_{257}$▶C, $T_{328}$▶G, $G_{329}$▶C, $T_{340}$▶G, $G_{341}$▶C) | HA | this study |
| **Flp-In T-REx-293- FAM136A knockout- FAM136A -Mock** | pcDNA5/FRT/TO and PB-CuO-MCSIRES-GFP-EF1-CymR-Puro | ORF *FAM136A* And -- | HA | this study |
| **Flp-In T-REx-293- FAM136A knockout- FAM136A - FAM136A** | pcDNA5/FRT/TO and PB-CuO-MCSIRES-GFP-EF1-CymR-Puro | ORF *FAM136A* And ORF *FAM136A* | HA and FLAG | this study |
| **Flp-In T-REx-293-MIA40** | pcDNA5/FRT/TO | ORF *MIA40* | Strep | (Petrungaro et al., 2015) |
| **Flp-In T-REx-293-MIA40-SPS** | pcDNA5/FRT/TO | ORF *MIA40* ($T_{157}$▶G; $T_{163}$▶G) | Strep | (Habich et al., 2019) |
| **Flp-In T-REx-293-MIA40-C53S** | pcDNA5/FRT/TO | ORF *MIA40* ($T_{157}$▶G) | Strep | (Fischer et al., 2013) |
| **Flp-In T-REx-293-MIA40-F68E** | pcDNA5/FRT/TO | ORF *MIA40* ($T_{202}$▶G, $T_{204}$▶G, $T_{203}$▶A) | Strep | (Habich et al., 2019) |
| **HAP1** | -- | -- | -- | (Carette et al., 2011) |
| **HAP1 DELE1 knockout** | pL-CRISPR.EFS.tRFP (Addgene 57819) | -- | -- | (Fessler et al., 2020) |
| **HAP1 HRI knockout** | pL-CRISPR.EFS.tRFP (Addgene 57819) | -- | -- | (Fessler et al., 2020) |
| **HEK293T** | -- | -- | -- | (Habich et al., 2019) |
| **YME1L knockout** | - | - | - | |
| **HEK293T-MIA40 knockdown #2** | pSpCas9(BB)2A-GFP | -- | -- | (Habich et al., 2019) |
| **HEK293T-MIA40 knockdown #2-Mock** | PB-CuO-MCSIRES-GFP-EF1-CymR-Puro | | -- | |
| **HEK293T-MIA40 knockdown #2-MIA40** | PB-CuO-MCSIRES-GFP-EF1-CymR-Puro | ORF *MIA40* | -- | |
| **HEK293T-AIFM1 knockout** | pSpCas9(BB)2A-GFP | -- | -- | (Salscheider et al., 2022) |
| **HEK293T-AIFM1 knockout-Mock** | PB-CuO-MCSIRES-GFP-EF1-CymR-Puro | | -- | (Salscheider et al., 2022) |
| **HEK293T-AIFM1 knockout-AIFM1** | PB-CuO-MCSIRES-GFP-EF1-CymR-Puro | ORF *AIFM1* | HA | (Salscheider et al., 2022) |
| **Rosetta2 (DE3)-MIA40** | pGEX-6p-1 | ORF MIA40 (C4▶S) | GST (N-terminal) | (Rothemann et al. 2024) |

| | | | | |
|---|---|---|---|---|
| **Rosetta2 (DE3)-MIA40 SPS** | pGEX-6p-1 | ORF MIA40 (C4►S; C53►S; C55►S) | GST (N-terminal) | (Rothemann et al. 2024) |
| **Rosetta2 (DE3)-MIA40 C53S** | pGEX-6p-1 | ORF MIA40 (C4►S; C53►S) | GST (N-terminal) | (Rothemann et al. 2024) |
| **Rosetta2 (DE3)-MIA40 F68E** | pGEX-6p-1 | ORF MIA40 (F68►E) | GST (N-terminal) | (Rothemann et al. 2024) |

#### Table S2. Primers and Plasmids

| Plasmid | Primer (5´-3´) | Restriction sites |
|---|---|---|
| **pMA-T-FAM136A-HA (gene synthesis: gene art)** | -- | NheI, KpnI, BamHI, HindIII, BstBI |
| **pMA-RQ-FAM136A-9CA-HA in (gene synthesis: gene art)** | -- | NheI, KpnI, BamHI, HindIII, BstBI |
| **pMA-RQ-FAM136A-CA-1-HA in (gene synthesis: gene art)** | -- | NheI, KpnI, BamHI, HindIII, BstBI |
| **pMA-RQ-FAM136A-CA-2-HA in (gene synthesis: gene art)** | -- | NheI, KpnI, BamHI, HindIII, BstBI |
| **pGEM4-FAM136A-M28A/L31A/M32A (gene synthesis: GenScript)** | - | KpnI, BamHI |
| **pGEM4-FAM136A-M28S/L31S/M32S (gene synthesis: GenScript)** | - | KpnI, BamHI |
| **FAM136A-HA in pCDNA5** | -- | KpnI, BamHI |
| **FAM136A -9CA-HA in pCDNA5** | -- | KpnI, BamHI |
| **FAM136A -CA-1-HA in pCDNA5** | -- | KpnI, BamHI |
| **FAM136A -CA-1-HA in pCDNA5** | -- | KpnI, BamHI |
| **FAM136A-FLAG in PB** | Fw: GCGGCTAGCGGTACCATCTCCATGGCTGAGCTG<br>Rev: CGCTTCGAAAAGCTTGGATCCTTACTTGTCGTCATCGTCTTTGTAGTCGCTTCC | NheI, BstBI |
| **Fam136A-HA in pGEM4** | -- | KpnI, BamHI |
| **Fam136A-9CA-HA in pGEM4** | -- | KpnI, BamHI |
| **FAM136A guide in pSpcas9(bb)-2a-GFP** | -- | Bbs1 |
| **FAM136A guide in pLentiCRISPRv2** | -- | Esp3I |
| **FAM136A guide in pL-CRISPR.EFs.GFP.P2A.Puro** | -- | Esp3I |

#### Table S3. Antibodies

| Antibody | Company | Identifier |
|---|---|---|
| **Rabbit polyclonal anti-FAM136A** | ThermoFisher Scientific | Cat# PA5-56345 |
| **Rabbit polyclonal anti-FAM136A** | Merck | Cat# HPA030104 |
| **Goat anti-Mouse lgG (H&L), HRP Conjugate** | ImmunoReagents | Cat# GtxMu-003-DHRPX |
| **Goat anti-Rabbit lgG (H&L), HRP Conjugate** | ImmunoReagents | Cat# GtxRb-003-DHRPX |
| **Mouse monoclonal anti-Actin** | ThermoFisher Scientific | Cat# #MA5-11869 |
| **Rabbit monoclonal anti-ATF4** | Cell Signaling Technology | Cat# #11815 |
| **Mouse monoclonal anti-CHOP** | Cell Signaling Technology | Cat# #2895 |
| **Rabbit polyclonal anti-CLPB** | Sigma-Aldrich | HPA039006 |
| **Mouse monoclonal anti-FLAG** | Sigma-Aldrich | Cat# F3165 |
| **Mouse monoclonal anti-GAPDH** | ProteinTech | Cat# 60004-1-Ig |
| **Rabbit polyclonal anti-GDF15** | ProteinTech | Cat# 27455-1-AP |
| **Mouse monoclonal anti-PDH** | Santa-Cruz | Cat# sc-377092; RRID:AB_2716767 |
| **Mouse monoclonal anti-PRDX2** | Sigma-Aldrich | WH0007001M1 |
| **Mouse monoclonal anti-TIM9** | Santa Cruz | Cat# sc-101284 |
| **Mouse Strep-tag antibody** | Qiagen | Cat# 34850 |
| **Rabbit Alexa Fluor 488** | Invitrogen | Cat# A11008 |
| **Rabbit polyclonal anti-AIF** | Merck | Cat# AB16501 |
| **Rabbit polyclonal anti-COX17** | Biorbyt | Cat# orb160552 |
| **Rabbit polyclonal anti-CPOX** | St John's Laboratory | Cat# STJ23214 |
| **Rabbit polyclonal anti-HA** | Sigma-Aldrich | Cat# SAB4300603; RRID:AB_10620829 |
| **Rabbit polyclonal anti-MIA40** | (Erdogan et al., 2018; Fischer et al., 2013) | N/A |
| **Rabbit polyclonal anti-MIC19** | Proteintech | Cat# 25625-1-AP |
| **Rabbit polyclonal anti-MIC25** | Proteintech | Cat# 20639-1-AP |
| **Rabbit polyclonal anti-NDUFA8** | Abcam | Cat# ab184952 |
| **Rabbit polyclonal anti-NDUFS5** | Abcam | Cat# ab179806 |
| **Rabbit polyclonal anti-TOM20** | Santa Cruz | Cat# sc-11415 |
| **Rabbit polyclonal anti-YME1** | Proteintech Europe | Cat# 11510-1-AP |
| **Rabbit polyclonal anti-HAX1** | St John's Laboratory | Cat# STJ27497 |

### Table S4. Oligonucleotides and siRNAs

| Oligonucleotides | Company | Identifier |
|---|---|---|
| **FAM136A Guide sgRNA2**<br>Fw: CACCGTCCGGAAGATGCAGGTAGCG | this study | N/A |
| **FAM136A Guide sgRNA2**<br>Rev: AAACCGCTACCTGCATCTTCCGGAC | this study | N/A |
| **FAM136A Guide sgRNAex1**<br>**Fw: CACCGACTCCACCGCCTCCTGCACC**<br>**Rev: AAACGGTGCAGGAGGCGGTGGAGTC** | this study | N/A |
| **FAM136A Guide sgRNAex2**<br>**Fw: CACCGCTGGTGCACCTGCTTCATGG**<br>**Rev: AAACCCATGAAGCAGGTGCACCAGC** | this study | N/A |
| **Non-targeting control sgRNA**<br>**Fw: CACCGGTATGTCGGGAACCTCTCC**<br>**Rev: AAACGGAGAGGTTCCCGACATACC** | this study | N/A |
| **DELE1 Guide sgRNA**<br>**Fw: CACCGAGCGACATGTGGCGCCTCCC**<br>**Rev: AAACGGGAGGCGCCACATGTCGCTC** | (Fessler et al., 2020) | N/A |
| **HRI Guide sgRNA**<br>**Fw: CACCGCCATCGACTTTCCCGCCGA**<br>**Rev: AAACTCGGCGGGAAAGTCGATGGC** | (Fessler et al., 2020) | N/A |
| **siRNA target sequence**<br>**TGCCAACATCCTGTTTACCAA** | Quiagen<br>SI04228301 | N/A |
| **siRNA target sequence**<br>**TACCAGACTCTTCTTACTACA** | Quiagen<br>SI04331789 | N/A |

#### Table S5. Further Tools and Equipment

| | Company/ Source | Identifier |
|---|---|---|
| **EasyTag™ Express Protein Labeling Mix, [35S]** | Perkin-Elmer | Cat# NEG772 |
| **EasyTag™ Express Protein Labeling Mix, [35S]** | Revvity (Germany) GmbH | NEG772002MC |
| **ROTI®Quant universal** | Carl Roth | Cat # 0120.1 |
| **TnT Quick Coupled Transcription/Translation System** | Promega | Cat# L1170 |
| **Pierce 660 nm Protein Assay Reagent** | Thermo Scientific | Cat# 22660 |
| **Carbonylcyanid-3-chlorphenylhydrazon (CCCP)** | Sigma-Aldrich | Cat# C2759 |
| **FuGENE HD Transfection Reagent** | Promega | Cat# E2311 |
| **RosettaTM 2 (DE3) SinglesTM Competent Cells** | Novagen | Cat# 70954-3 |
| **One Shot TOP10 Chemically Competent E. coli** | Thermo Fisher | Cat# C404010 |
| **TMRM** | TCI chemicals | Cat# 115532-50-8 |

**Software and Algorithms**

| | | |
|---|---|---|
| **Alpha-fold Multimer** | (Jumper et al., 2021) | |
| **Pymol** | | https://www.pymol.org |
| **Fiji** | (Schindelin et al., 2012) | https://imagej.net/Fiji |
| **Image Lab 5.2** | Biorad Laboratories | |
| **ImageQuant TL 8.1** | GE Healthcare Life Sciences | |
| **iMTS score** | (Boos et al., 2018) | (http://iomiqsweb1.bio.uni-kl.de/). |
| **Graphpad Prism** | GraphPad Software, Boston, Massachusetts USA | www.graphpad.com |

#### SUPPLEMENTAL FIGURES

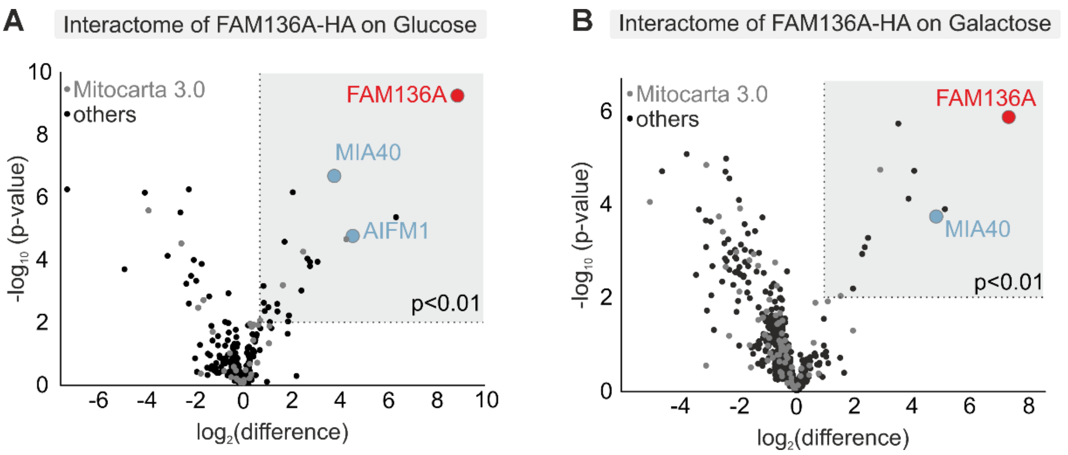


**Supplemental Figure S1. FAM136A interacts with MIA40 and AIFM1.**

**A., B.** Interactome of FAM136A-HA from cells grown on glucose (A) or galactose (B) as carbon sources. Cells were treated with NEM to avoid aberrant disulfide formation, and lysed under native conditions. FAM136A-HA was enriched using HA-antibody beads and eluates were analyzed by label-free mass spectrometry. The experiment relies on the analysis of 4 biological replicates. Fold enrichment in the FAM136A-HA IP was plotted for significant hits. N = 4 biological replicates, an unpaired one-sample two-sided Student's t-test was applied ($p<0.01$, log2-enrichment >1).

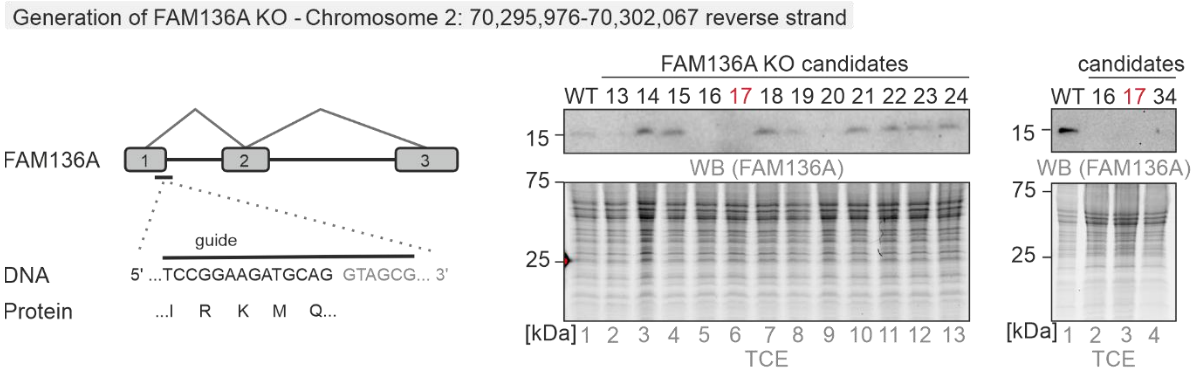


**Supplemental Figure S2. Generation of FAM136A knockout cells using CRISPR-Cas technology.**

A guide directed against the first exon of FAM136A gave rise to multiple clones. Successful targeting of the gene was confirmed by immunoblotting against FAM136A.

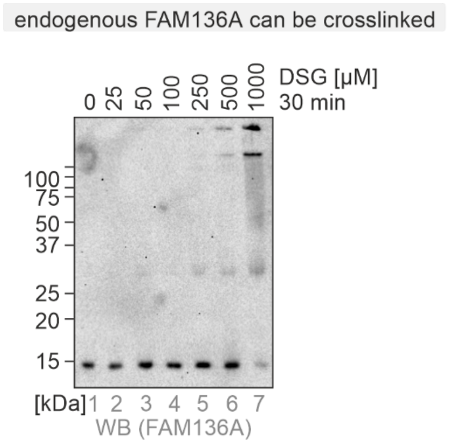


**Supplemental Figure S3. Endogenous FAM136A can be crosslinked.**

HEK293 cells were incubated for 30 min with the indicated amounts of the crosslinker disuccinimidylglutarat (DSG). The crosslinking reaction was stopped by addition of TRIS. Cells were lysed in loading buffer containing SDS and DTT, and analysed by subsequent immunoblotting against FAM136A. FAM136A can be partially crosslinked to a band of about 30-35 kDa in size, and at higher DSG concentrations also to higher MW oligomers.

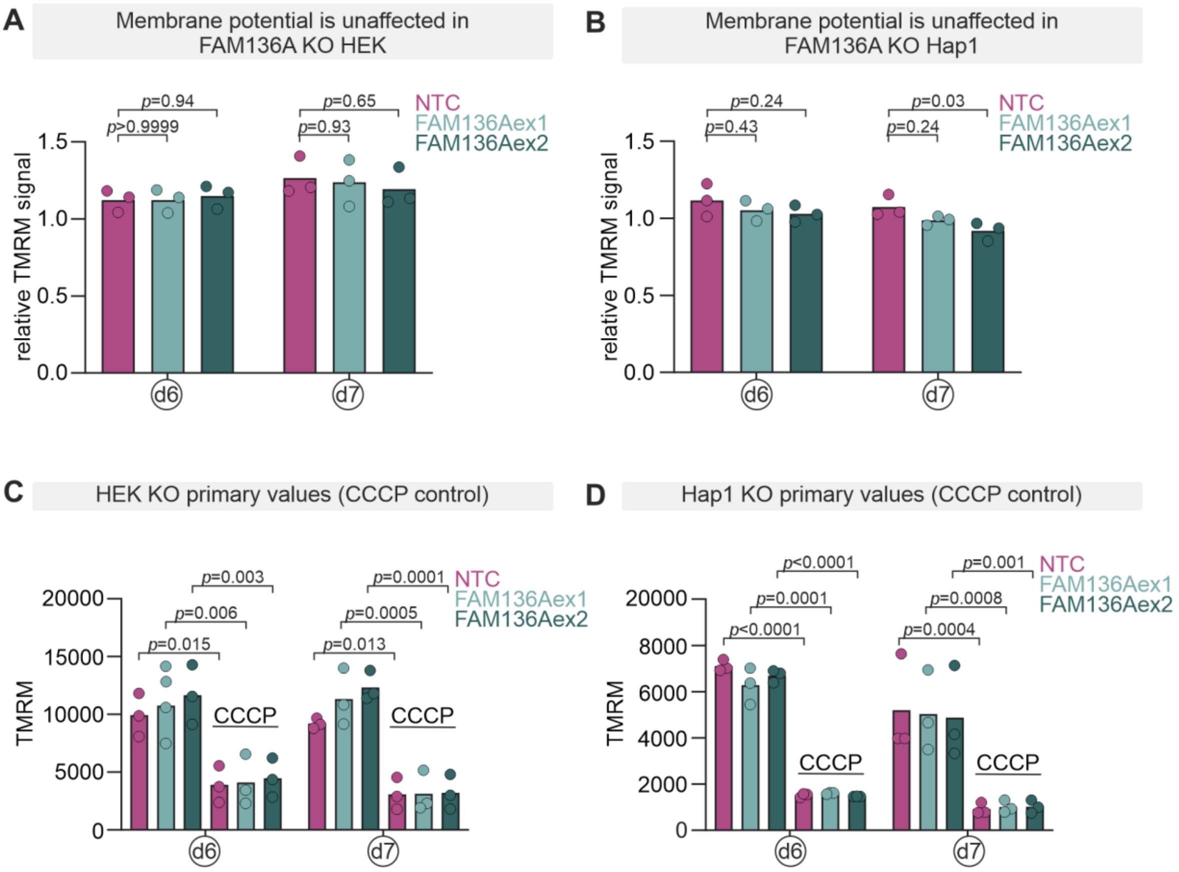


**Supplemental Figure S4. Mitochondrial membrane potential upon acute loss of FAM136A.**

**A., B.** HEK293T (A) or HAP1 (B) cells transduced with sgRNAs against FAM136A exon 1, exon 2 or a non-targeting sgNTC were incubated for 30 min with 100 µM TMRM +/- 20 µM CCCP. Cells were measured by flow cytometry and TMRM fluorescence of sgRNA transduced cells was normalised to non-transduced cells from the same well. Two-way ANOVA.

**C., D.** Primary TMRM values of the transduced cells for comparison with the CCCP positive control. Two-way ANOVA.

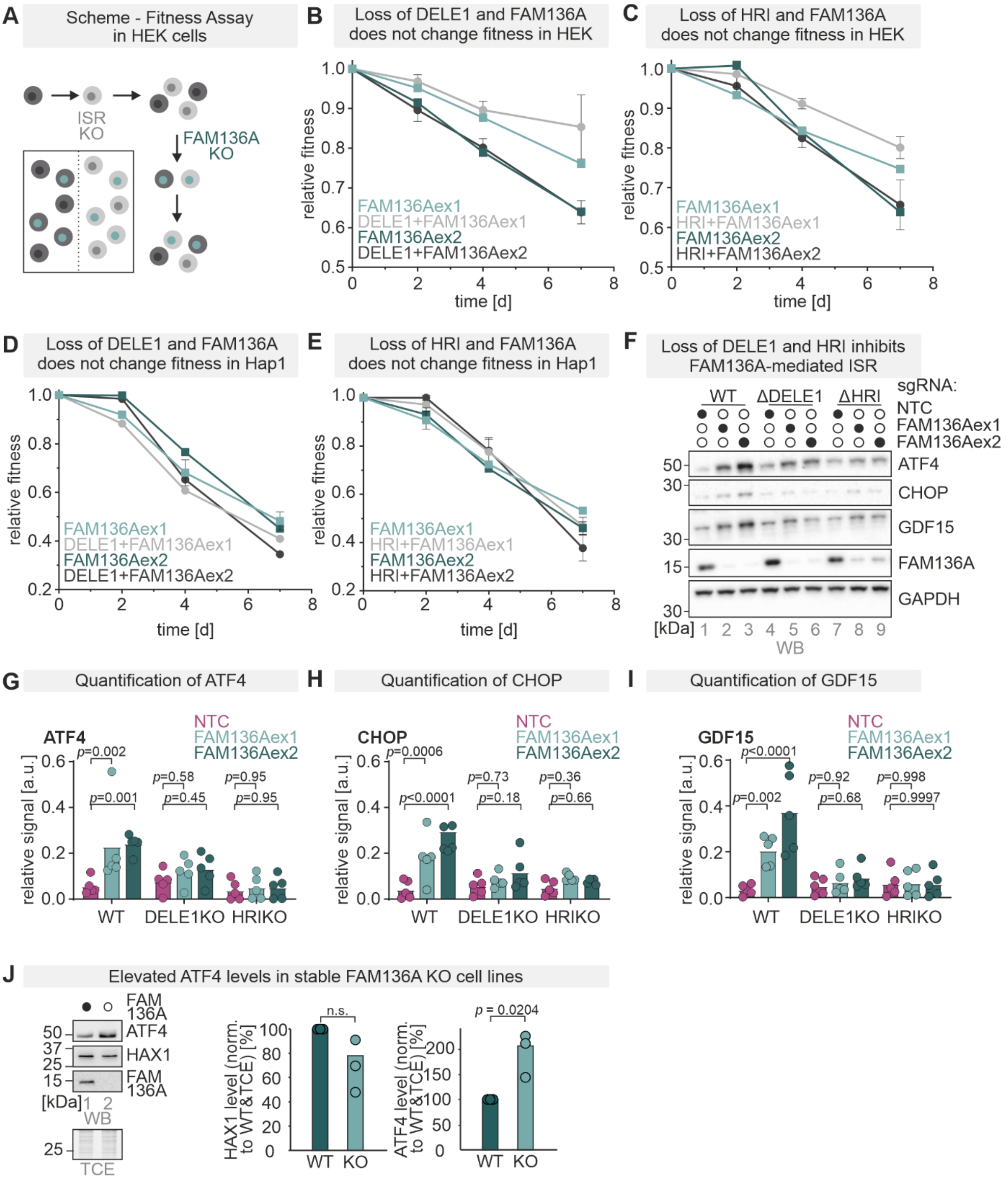


**Supplemental Figure S5. The ISR is not responsible for loss of cellular fitness upon FAM136A loss**

**A.-E.** Fitness assay in DELE1/HRI deficient cells. A competitive fitness assay (A) was performed in HEK293T (B., C.) or HAP1 (D., E.) cells. They were transduced with a non-fluorescent non-targeting guide (sgNTC) or GFP-containing guides against DELE1/HRI. sgNTC cells were mixed with sgDELE1/sgHRI respectively and transduced with mScarlet-containing sgNTC or sgRNAs against FAM136A exon 1 or exon 2. The fraction of mScarlet$^{+}$ cells was measured over time and analysed separately for sgNTC or sgDELE1/HRI (GFP$^{+}$) cells from the same well.

**F.** HAP1 wildtype or DELE1/HRI deficient cells were transduced with two different sgRNAs to ablate FAM136A. After 6- or 7-days cells were lysed and analysed by immunoblotting for the indicated proteins.

**G.-I.** Quantification of proteins shown in F. N = 5 replicates. Two-way ANOVA.

**J.** ATF4 levels are elevated in chronic FAM136A KO cells (n=3) whereas HAX1 levels are not affected by chronic loss of FAM136A (n=3) one-sample t-test.

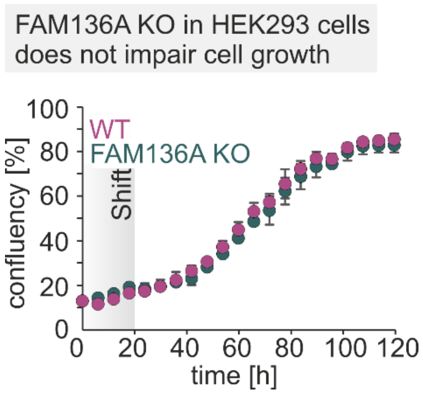


**Supplemental Figure S6. Cells with chronic loss of FAM136A adapt**

CRISPR/Cas9 FAM136A KO cells do not differ in cell growth from WT cells after exchanging the carbon source after 20h to galactose-containing medium. Cell confluency was measured every 6 hours using Cytosmart OMNI (n=4).
